# Small intestine microbiota development prevents early-life adiposity via IL-22-mediated intestinal PPARα suppression

**DOI:** 10.64898/2026.06.11.731695

**Authors:** Catherine D. Shelton, Camila Bernardo de Brito, Vinícius Dias Nirello, Nicole Kirchoff, Julia Lane, Joanna Olivas, Darian T. Carroll, Derek Armstrong, Matthew W. Rhee, Merrygay N. James, Louise Lantier, Marco Aurélio Ramirez Vinolo, Mariana Xavier Byndloss

## Abstract

Perturbation to the early-life microbiota has long-term detrimental effects on health and development, leading to increased risk for metabolic dysfunction and childhood obesity. Despite the central role of the small intestine (SI) in energy balance, the impact of SI microbiota establishment on the regulation of host metabolism and early-life adiposity remains unclear. Here, we report that disruption of a critical SI microbiota-intestinal epithelial cell circuit, specifically during a critical early-life period, drives long-lasting obesity. We demonstrate that the SI microbiota expands in abundance and diversity significantly between 2 and 3 weeks of life, and that segmented filamentous bacteria (SFB) and *Lactobacillus intestinalis* establish residence. Disruption of the early-life SI microbiota with antibiotics leads to enhanced lipid uptake and adiposity, driven by increased peroxisome proliferator-activated receptor alpha (PPARα) expression and activity in SI epithelial cells (IECs). We demonstrate that SFB and *L. intestinalis* are key regulators of PPARα in SI IECs by increasing intestinal IL-22 levels specifically during weaning, which is necessary for inhibition of antibiotic-induced adiposity in a PPARα-dependent manner. Together, this work provides mechanistic insights into beneficial microbiota-induced epithelial-immune crosstalk in the SI that is specific to early life, a critical protective mechanism against excessive adiposity in infancy, and offers insight into how antibiotics during infancy may increase the risk of childhood obesity.

**GRAPHICAL ABSTRACT:** 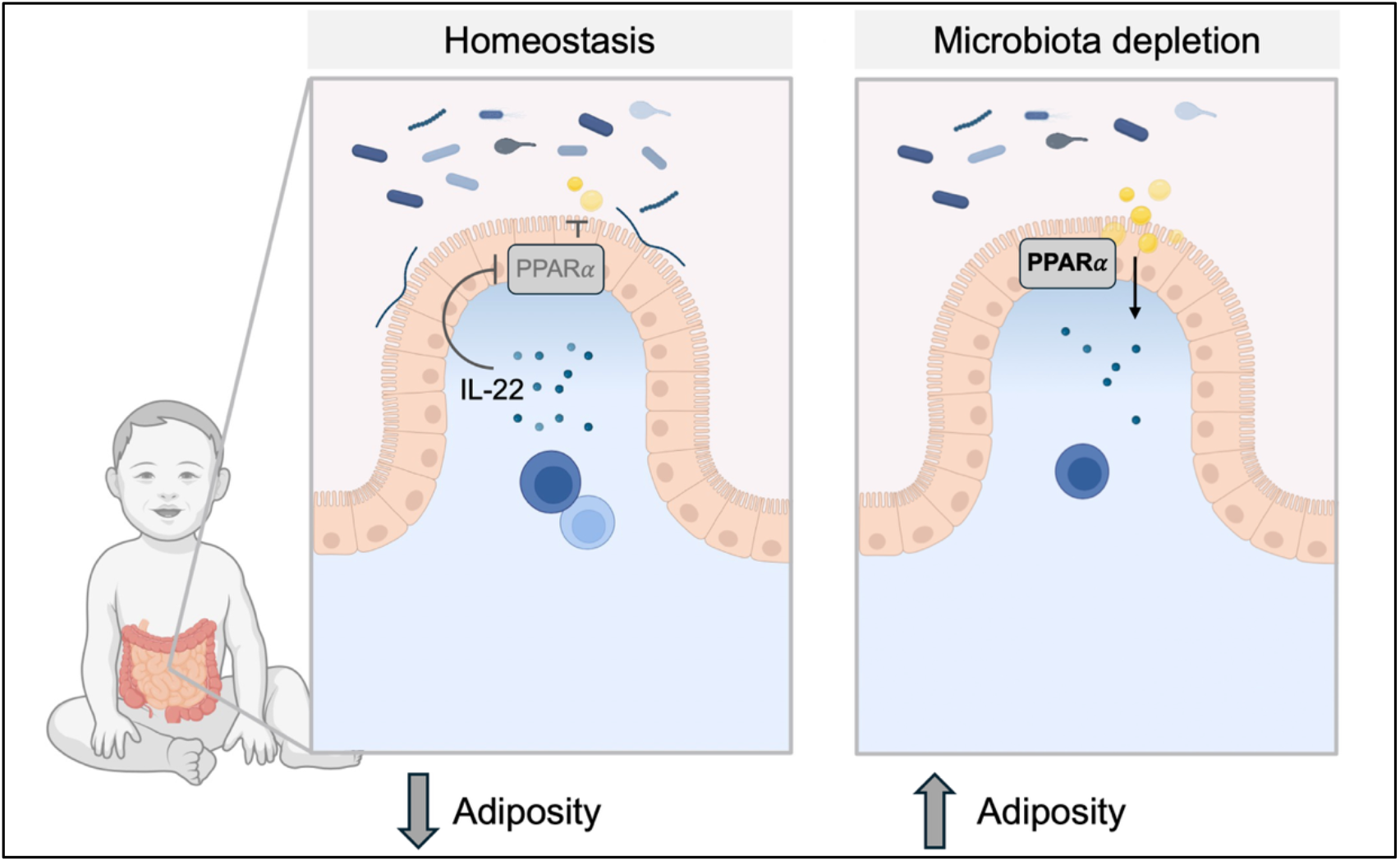

## INTRODUCTION

The first 1,000 days of life, from conception until a child is two years old, are a critical window for establishing long-term health (*1*). Perturbations to a child’s normal growth and development during this timeframe result in sustained effects (*1*). For instance, early-life environmental factors, such as antibiotic treatment, have been shown to increase an infant’s risk for developing childhood obesity (*2-6*). Multiple studies have demonstrated that treatment with antibiotics, such as penicillin and macrolides, during a child’s first two years of life is associated with higher body mass index (BMI) and greater adiposity in adolescence (*7-11*). Despite substantial evidence linking early-life antibiotic treatment and obesity, the mechanisms by which antibiotic treatment during infancy impacts host metabolism remain unknown.

Previous studies suggest that antibiotic treatment may increase the risk of childhood obesity by disrupting the early-life gut microbiota, a highly dynamic microbial community that matures during infancy (*12-14*). Establishment of the gut microbiota promotes immune system development and inhibits enteric pathogen colonization (*15, 16*), but the mechanisms by which the early-life gut microbiota regulates host metabolism remain poorly understood. In mouse models, changes to the gut microbiota induced by early-life exposure to low doses of penicillin (LDP) were sufficient to increase adiposity when the LDP-treated microbiota was transferred to gnotobiotic mice (*17*). However, translating these insights from mice to children requires a clearer understanding of the mechanisms linking the early-life microbiota to host adiposity.

Characterization of the infant gut microbiota is predominantly based on data derived from 16S rRNA sequencing of fecal samples (*12, 13*). Nevertheless, the microbiota is not homogeneous throughout the gastrointestinal tract; instead, different regions of the intestine harbor distinct microbial communities that serve specific functions (*18, 19*). For instance, the small intestine (SI) microbiota, which is less dense and diverse than the fecal microbiota, plays a significant role in regulating host adiposity and metabolism (*20, 21*). Members of the SI microbiota regulate small intestine epithelial cells (IECs), which are responsible for the majority of nutrient absorption (*20-22*). Although previous studies in adult mice have identified the protective role of the SI microbiota in modulating nutrient uptake by IECs in models of diet-induced obesity (DIO) (*23, 24*), the contribution of the early-life SI microbial community to the regulation of nutrient uptake during infancy remains unknown. Importantly, the development of the infant SI microbiota is understudied, as characterization of the early-life SI microbiota is largely based on samples from children and infants with SI disease (*25*). Moreover, the early-life SI epithelium differs from that of an adult, in both cell-type composition and transcriptional profiles, and matures as an infant develops (*26-29*). Therefore, interactions discovered between the adult SI microbiota and IECs during high-fat (HF) diet exposure are likely distinct from microbiota-IEC crosstalk in the early-life intestine, especially as members of the adult SI microbiota may not be present in the infant SI microbial community. As early life is a significant developmental period, infant SI microbiota-mediated regulation of intestinal nutrient uptake may be a key component in establishing long-lasting metabolic health, with disruption to these critical interactions leading to sustained perturbation in host metabolism (*1*). To understand how the infant SI microbiota protects against metabolic dysfunction, we used a mouse model of early-life disruption of the SI microbial community to characterize its development during infancy and investigate how expansion of specific SI microbiota members promotes pivotal beneficial IEC-immune interactions that control intestinal lipid uptake and adiposity.

## RESULTS

### The small intestine microbiota rapidly expands and increases in diversity during early life

In contrast to humans, intestinal development in mice occurs after birth, and the mouse intestine only acquires characteristics of a mature SI (e.g., crypts and villi) at 14 days of age (*30*). Thus, we began our investigation into early-life microbiota-SI interactions at 2 weeks of age, an age when the mouse intestine begins to resemble the infant intestine in humans (*31, 32*). As the early-life SI microbiota remains largely uncharacterized, we first measured the total bacterial burden in the distal SI of 2- and 3-week-old mice (**Fig. 1A**). We determined that the SI microbiota significantly increased in abundance between 2 and 3 weeks of age, a time period in which mice are weaning from milk and transitioning to eating solid food (**Fig. 1A**) (*33*). In contrast, there was no significant difference in bacterial abundance in the colon between 2- and 3-week-old mice (**Fig. 1A**), suggesting that this early-life bacterial expansion is specific to the SI microbiota. Metagenomic sequencing revealed that the SI microbiota of 2- and 3-week-old mice differed significantly, both in terms of genus composition and the pathways present (**Fig. 1, B – D**). Only two bacterial species significantly increased in the SI between 2 and 3 weeks, Segmented Filamentous Bacteria (SFB, denoted as Candidatus Arthromitus in **Fig. 1B**) and *Lactobacillus intestinalis* (**Fig. 1, E and F**). To interrogate the impact of SI microbiota development on the host, we utilized an established model of microbiota perturbation in which low doses of penicillin (LDP) were given in the drinking water from birth until mice were 3 weeks old (**Fig. 1G**) (*17, 23*). Exposure to early-life antibiotics did not alter the SI microbiota after 2 weeks of exposure (data not shown) but did significantly alter the composition of the SI microbiota after 3 weeks of treatment (**Fig. 1H**) and reduced the abundance of SFB and *L. intestinalis* (**Fig. 1, I and J**). Antibiotic treatment affected the colon microbiota at both 2 and 3 weeks (**Fig. S1, A – C**) but did not significantly affect its expansion and development (**Fig. 1A** and **Fig. S1, D - F**). For instance, only *Streptococcus*, a minor member of the colonic microbiota, was both significantly different in the colon between 2 and 3-weeks in mock mice as *Streptococcus* abundance decreased in the colon of 3-week-old mice compared to 2-week-old mice and was depleted in the colon of 3-week-old LDP-treated mice **(Fig. S1, E** and **F)**. Our results demonstrate that the distal SI microbiota undergoes a marked expansion during the weaning period, characterized by increased abundances of SFB and *L. intestinalis*, and that early-life antibiotics impair this developmental process.

**Figure 1.**
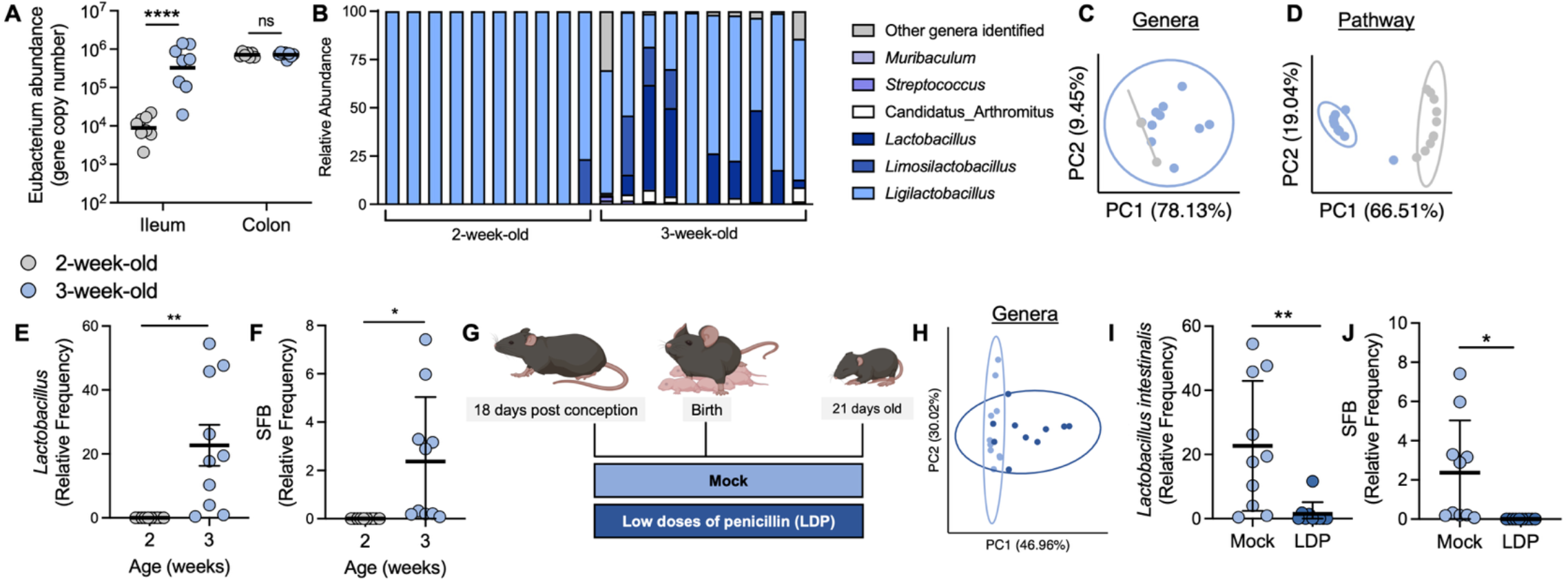
Early-life antibiotics impair the development of the small intestine (SI) microbiota. (**A**) Total bacterial abundance in the distal SI of 2- and 3-week-old conventional mice as determined by 16S rRNA qPCR. (**B**) Relative abundance of genera in the SI of mock mice at 2 and 3 weeks of age as determined by metagenomic sequencing. (**C and D**) PCoA plot of (**C**) genera and (**D**) pathways in 2 and 3-week-old SI content based on Bray-curtis dissimilarity matrix. Statistical significance was determined by PERMANOVA (C and D) p = 0.001. (**E and F**) Relative frequency of (**E**) *Lactobacillus intestinalis* and (**F**) segmented filamentous bacteria (SFB) in 2 and 3-week-old SI content. (**G**) Mouse model of early-life antibiotic treatment. (**H**) PCoA plot based on Bray-curtis dissimilarity of the genera in 3-week-old Mock and LDP SI content. Statistical significance was determined by PERMANOVA, p = 0.026. (**I and J**) Relative frequency of (**I**) *Lactobacillus intestinalis* and (**J**) segmented filamentous bacteria (SFB) in 3-week-old SI content from mock and antibiotic treated mice. (A, C – F, H - J) Each dot represents one animal. (B) Each bar represents one animal. (A) Bars represent geometric mean. (C, D, and H) Circles represent 95% confidence interval. (E, F, I, J) Bars represent mean ± standard error of the mean. (A) ****, p < 0.0001 using a two-way anova with Šídák’s multiple comparisons test. (E, F, I, J) *, p < 0.05; **, p < 0.01 using an unpaired two-tailed Welch’s t test.

### Exposure to early-life antibiotics alters small intestinal absorptive cells and increases lipid uptake

We next investigated how changes in the early-life SI microbiota impact intestinal epithelial development. Between 2 and 3 weeks of life, the SI undergoes substantial maturation, as crypt length increases and villi shorten (**Fig. 2A**), but how the SI microbiota impacts this transition is largely unknown (*29*). Therefore, we performed single-cell RNA sequencing (scRNA-seq) of crypts isolated from 3-week-old mock and LDP-treated mice. Antibiotic treatment significantly altered the composition of SI IEC populations, promoting the expansion of proliferative cells (stem cells and transit-amplifying cells) and tuft cells, while reducing the abundance of goblet cells (**Fig. 2B**). Further characterization of the intestinal epithelium supported scRNA-seq results as antibiotic-exposed mice had a significant increase in proliferating cells, as demonstrated by increased EdU and Ki67 staining in the crypts, compared to mock mice (**Fig. S2, A – D**). Reductions in UEA-1 (a goblet cell marker) staining in LDP-treated mice also corroborate the scRNA-seq results (**Fig. S3, A and B**). In addition, crypts isolated from LDP-treated mice formed significantly more enteroids than crypts from 3-week-old mock mice (**Fig. S2A**), suggesting an increased proliferative capacity of the intestinal epithelium. Together, these data indicate that early-life disruption of the SI microbiota significantly affects the composition and function of the SI epithelium.

**Figure 2.**
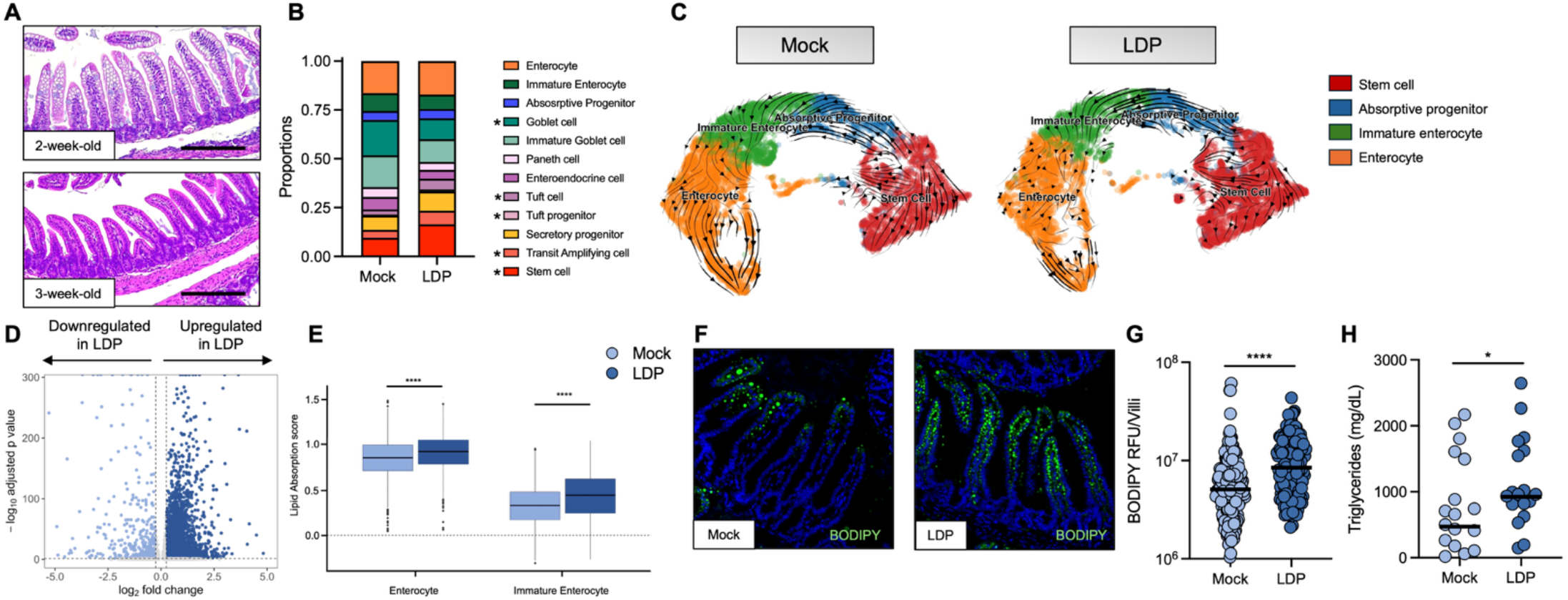
Early-life antibiotics perturb the intestinal epithelium and increase lipid absorption. **(A)** Representative images of the distal small intestine (SI) from 2 and 3-week-old mice. Scale bar represents 200 µm. (**B**) Proportion of the defined populations for broad epithelial classification in SI crypts isolated from 3-week-old Mock and LDP crypts. (**C**) RNA velocity of absorptive epithelial cell population in Mock and LDP samples calculated using *scVelo*. (**D**) Volcano plot of differentially expressed genes in the absorption epithelial cell population in LDP cells compared to Mock. (**E**) Lipid absorption score in enterocytes and immature enterocytes in Mock and LDP-treated mice. (**F - H**) 3-week-old Mock and LDP exposed mice were fasted for 4 hours and then challenged with a bolus of olive oil containing the fluorescent fatty acid BODIPY-C12. (**F**) Representative immunofluorescent images of ileum sections 3 hours after fat challenge from Mock and LDP mice. (**G**) Quantification of BODIPY-C12 fluorescent intensity in the intestine. (**H**) Serum triglycerides of Mock and LDP-exposed mice 3 hours after olive oil challenge. (B) *, p < 0.05 using Speckle. (D) Genes were considered significantly differentially expressed using a Wilcoxon rank-sum test with Bonferroni-adjusted p < 0.05 and |log2FC| > 0.25. (E) The center line of the box plot represents the mean, the box represents the 25th–75th percentile, and the whiskers are 5th–95th percentile. ****, p < 0.0001 using a Wilcoxon test. (G) Dots represent individual villi, and the line represents geometric mean. (H) Each dot represents an individual mouse. (G) ****, p < 0.0001 using an unpaired Student’s t test. (H) *, p < 0.05 using a one-tailed Student’s t test.

We next focused our SI epithelium scRNA-seq analysis on absorptive-lineage cells, as previous work in adult DIO mouse models determined that SI microbiota members altered lipid uptake in IECs (*20, 22-24*). RNA velocity analysis indicated changes in the differentiation of the absorptive cells, with cells from antibiotic-treated mice shifted toward earlier velocity pseudotime states compared with mock mice (**Fig. 2C**). In line with this, over 2,000 genes were upregulated in absorptive cells from antibiotic-treated mice, suggesting substantial perturbation to the transcriptome of these cells due to early-life LDP exposure (**Fig. 2D**). Nutrient absorption genes, specifically those involved in lipid uptake, were significantly upregulated in mature and immature enterocytes from young mice exposed to antibiotics (**Fig. 2F**) (*34*). As excessive lipid uptake can promote host metabolic dysfunction (*35*), we next tested the ability of enterocytes to absorb lipids in our model. To this end, 3-week-old mock and LDP-treated mice were challenged with a bolus of olive oil containing BODIPY-C12. Immunofluorescent imaging revealed that antibiotic-exposed mice absorbed significantly more BODIPY-C12 than mock-treated mice (**Fig. 2, F and G**), thus validating our scRNAseq findings. In addition, LDP-treated mice had higher serum triglyceride levels after a fat challenge, supporting the hypothesis that antibiotic exposure increases lipid absorption (**Fig. 2H**). These findings indicate that LDP treatment during early life alters the nutrient-uptake capacity of absorptive IECs, leading to increased lipid uptake.

### Depletion of early-life small intestine microbiota members promotes sustained increases in adiposity

As early-life antibiotics promoted greater lipid absorption in young mice, we next investigated whether LDP treatment increased adiposity. Before the expansion of the SI microbiota (**Fig. 1A**), when mice were 2 weeks old, no difference was observed in the amount of abdominal fat accumulated between antibiotic- and mock-exposed mice (**Fig. 3A**). However, by 3 weeks of age, both male and female LDP-treated mice possessed significantly more abdominal fat than mock-treated mice (**Fig. 3B**). Antibiotic exposure from only 2 to 3 weeks of life was sufficient to increase adiposity in male mice, isolating the growth-promoting effects of antibiotics to this specific period of time (**Fig. 3C**) which coincides with the expansion of the SI microbiota (**Fig. 1A**). Body composition analysis indicated that LDP exposure promoted greater fat and lean mass in 3-week-old mice but did not alter food consumption (**Fig. S4, A - C**). Additional analysis using metabolic cages revealed no difference in the respiratory exchange ratio of mock and antibiotic-treated mice (**Fig. S4D**). Interestingly, no significant difference in energy expenditure between mice exposed to LDP and mock-treated mice was observed when body mass was considered (**Fig. S4, E -G**). Together, our data demonstrate that early-life exposure to antibiotics increases adiposity independently of food intake and energy expenditure, suggesting an alternative mechanism by which antibiotic-mediated disruption of the SI microbiota promotes adiposity.

**Figure 3.**
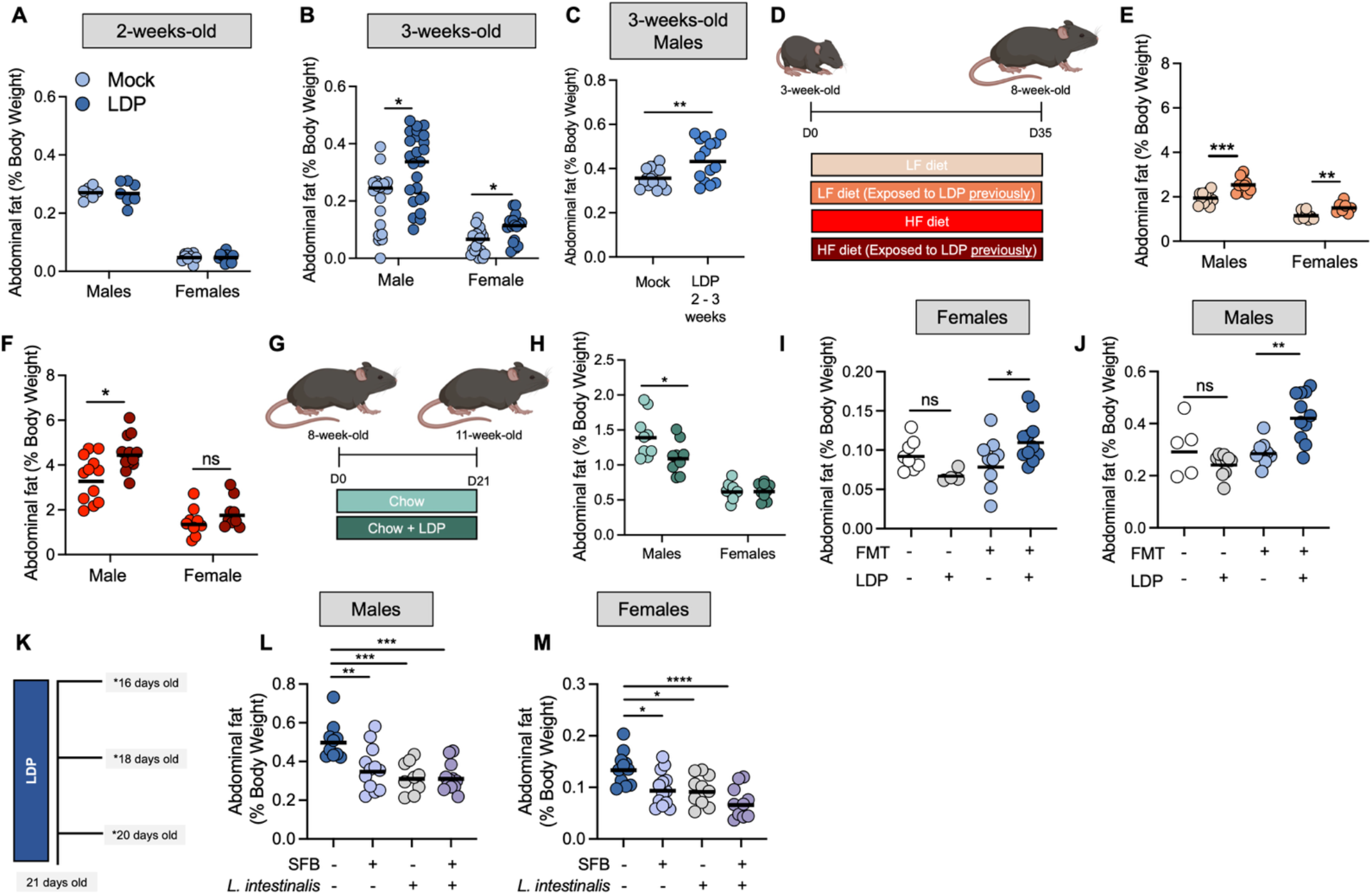
Disruption to the early-life small intestine microbiota increases adiposity. **(A and B)** Abdominal fat (% total body weight) of (**A**) 2-week-old and (**B**) 3-week-old mice mock treated or exposed to low dose penicillin (LDP). (**C**) Abdominal fat (% total body weight) of 3-week-old mice given low dose penicillin from 2 – 3 weeks and mock treated mice. (**D**) Diagram of long-term mouse experiment and groups. (**E and F**) Abdominal fat (% total body weight) of mice Mock treated or given LDP from birth until 3 weeks of age and then placed on a (E) low fat (LF) diet or (F) high fat (HF) diet for five weeks. (**G**) Model of adult LDP exposure. (**H**) Abdominal fat (% total body weight) of 8-week-old mice exposed to LDP for 3 weeks or mock-treated. (**I and J**) Germ-free (GF) breeders were either gavaged with a fecal microbiota transplant (FMT) or left GF. Breeders were then exposed to low doses of penicillin or mock-treated until their offspring were 3 weeks old. Abdominal fat (% total body weight) of 3-week-old (**I**) females and (**J**) males was then measured. (**K**) Experimental timeline for data shown in (**L** and **M**). Stars indicate days in which mice received microbes. (**L and M**) Abdominal fat (% total body weight) of (**L**) male or (**M**) female 3-week-old mice gavaged every 48 hours with SFB, *L. intestinalis*, both, or vehicle starting at postnatal day 16. Each dot represents one mouse. (B, E, F, and H) *, p < 0.05; **, p < 0.01; ***, p < 0.001 using a two-way anova with Šídák’s multiple comparisons test. (C) **, p < 0.01 using an unpaired Welch’s t test. (I, J, L, and M) *, p < 0.05; **, p < 0.01; ***, p < 0.001; ****, p < 0.0001 using a one-way anova with Dunnett’s multiple comparisons test.

As antibiotic exposure in the first two years of life is correlated with a greater risk of childhood obesity (*7, 10, 36*), we next asked whether LDP treatment resulted in long-lasting changes in host adiposity. We repeated our model in which mice are exposed to LDP (or mock-treated) for the first 3 weeks of life (**Fig. 1G**) and then, at weaning, we placed the mice on a low-fat (LF) or high-fat (HF) diet for 5 weeks until they reached adulthood at 8-weeks-old (**Fig. 3D**) (*37*). Despite no longer receiving antibiotics, mice exposed to LDP during infancy possessed greater abdominal fat than mock mice when fed a LF diet (**Fig. 3E**). When mice were weaned onto a HF diet, only male mice that received LDP during infancy had increased fat accumulation than mock-treated mice (**Fig. 3F**). While female mice given antibiotics had a slight increase in fat compared to mock mice, there was no significant difference (**Fig. 2F**). Having determined that early-life treatment with LDP resulted in sustained increases in fat accumulation, we next asked whether antibiotic exposure needed to occur during infancy to promote excess adiposity. Adult mice (8-week-old) were given LDP in their drinking water for 3 weeks (same duration as early-life treatment) (**Fig. 3G**). In contrast to young mice, antibiotic exposure reduced abdominal fat in 11-week-old male mice and did not change adiposity in 11-week-old female mice (**Fig. 3H**). Together, these data establish that antibiotic exposure during a specific, early-life window promotes long-lasting excess adiposity.

As LDP disrupted the development of the SI microbiota, we interrogated whether the increased adiposity seen in mice exposed to early-life antibiotics was dependent on perturbation to the gut microbiota. Gnotobiotic breeders were either left germ-free (GF) or gavaged with a fecal microbiota transplant (FMT) prepared from intestinal contents of the distal SI, cecum, and colon of conventional mice. Then, the same model performed previously (**Fig. 1G**) was applied using GF or conventionalized breeders, and LDP was added to the drinking water until their offspring were 3 weeks old. In contrast to offspring of breeders given an FMT, GF mice exposed to early-life antibiotics did not gain more abdominal fat than mock-treated GF mice (**Fig. 3, I and J**). As antibiotics did not increase adiposity in young GF mice, we next asked whether the specific members of the early-life SI microbiota depleted by antibiotics (SFB and *L. intestinalis*) (**Fig. 1, I and J**) protected young mice from fat accumulation. SFB, *L. intestinalis*, or both microbes were directly reintroduced into the microbiota of conventional LDP young mice via oral gavage every 48 hours from postnatal day 16 to 21 (**Fig. 3K**). The microbes were not reintroduced into the microbiota of the dams, which enabled us to test the direct effects of SFB and *L. intestinalis* on the infant mice physiology independently of their abundance in the maternal microbiota. Both SFB and *L. intestinalis* reduced abdominal fat in male and female 3-week-old mice, with a stronger effect observed when the microbes were given in combination (**Fig. 3, L and M**). We determined that depletion of SI microbiota members, specifically during early life, promotes adiposity.

### Early-life antibiotics disrupt microbiota-mediated downregulation of intestinal PPARα, leading to increased adiposity

Although we discovered that SI microbiota members regulated adiposity in young mice, the mechanism by which the early-life SI microbiota altered IEC nutrient uptake and host physiology remained unknown. To address this gap in knowledge, we returned to our scRNA-seq intestinal epithelium dataset (**Fig. 2B and C**). Candidate genes were prioritized by intersecting differentially expressed genes between mock and antibiotic treated groups across absorptive-lineage populations, including stem cells, absorptive progenitors, immature enterocytes and mature enterocytes, with genes showing differential transcriptional kinetics inferred by RNA velocity analysis. This identified 14 genes that were upregulated in antibiotic-treated mice and were supported by altered transcriptional dynamics (**Fig. 4A**). Among these, we selected *Ppara* for further investigation (**Fig. 4A**) as it encodes PPARα (peroxisome proliferator-activated receptor-α), a transcription factor that plays a key role in lipid metabolism (*38, 39*). RNA velocity analysis revealed that, unlike mock-treated mice, where *Ppara* showed limited variation across velocity pseudotime, antibiotic-exposed mice displayed clear trajectory-associated changes in *Ppara* expression (**Fig 4B**). The higher expression, particularly in proliferative and immature absorptive populations, suggests that microbiota regulation of *Ppara* may play a key role in dictating the differentiation and function of IECs in the absorptive lineage (**Fig. 4B**). Previous work determined that PPARα in IECs is required to increase lipid absorption, causing excess adiposity during consumption of an HF diet in adult mice (*40*). However, the contribution of intestinal PPARα to early-life excessive adiposity, in the absence of HF diet exposure, has not been characterized. Therefore, we sought to understand the role of microbiota-mediated regulation of PPARα in the SI during early life. *Ppara* was consistently upregulated in antibiotic-treated mice across all absorptive lineage populations (**Fig. 4A**). As PPARα functions as a transcriptional regulator, we also analyzed expression of PPARα target genes in our scRNA-seq intestinal epithelial dataset (**Fig. S5A**). Target genes of PPARα related to lipid uptake were upregulated in absorptive progenitor cells, immature enterocytes, and mature enterocytes of LDP-treated young mice, with particular enrichment of genes involved in lipid absorption and utilization in mature enterocytes (**Fig. S5A**). In addition to scRNAseq, we confirmed that expression of *Ppara* and its target genes related to lipid uptake and accumulation (*Fatp1, Fatp2*, and *Plin2*) were elevated in bulk SI epithelial cells from 3-week-old LDP-treated mice (**Fig. 4C**) (*40*). Consistent with our hypothesis that expansion of the SI microbiota regulates *Ppara* expression in IECs, no differences in the expression of *Ppara* and its target genes were observed in 2-week-old mock and LDP-exposed SI epithelial cells, an age at which no differences were observed in the SI microbiota (data not shown) (**Fig. S5B**). Similarly, no difference was seen in PPARα abundance in the colon or in the expression of *Ppara* in the liver or adipose tissue of 3-week-old mock and antibiotic-treated mice (**Fig. S5, C and D**), providing further support that exposure to LDP specifically perturbs *Ppara*-SI microbiota interactions during early life. Significantly, 3-week-old male PPARα knockout (KO) mice failed to exhibit increased adiposity induced by LDP exposure, whereas their heterozygous littermate controls accumulated more fat following LDP treatment (**Fig. 4D**). These data suggest that disruption to the early-life SI microbiota upregulates intestinal *Ppara*, resulting in increased adiposity.

**Figure 4.**
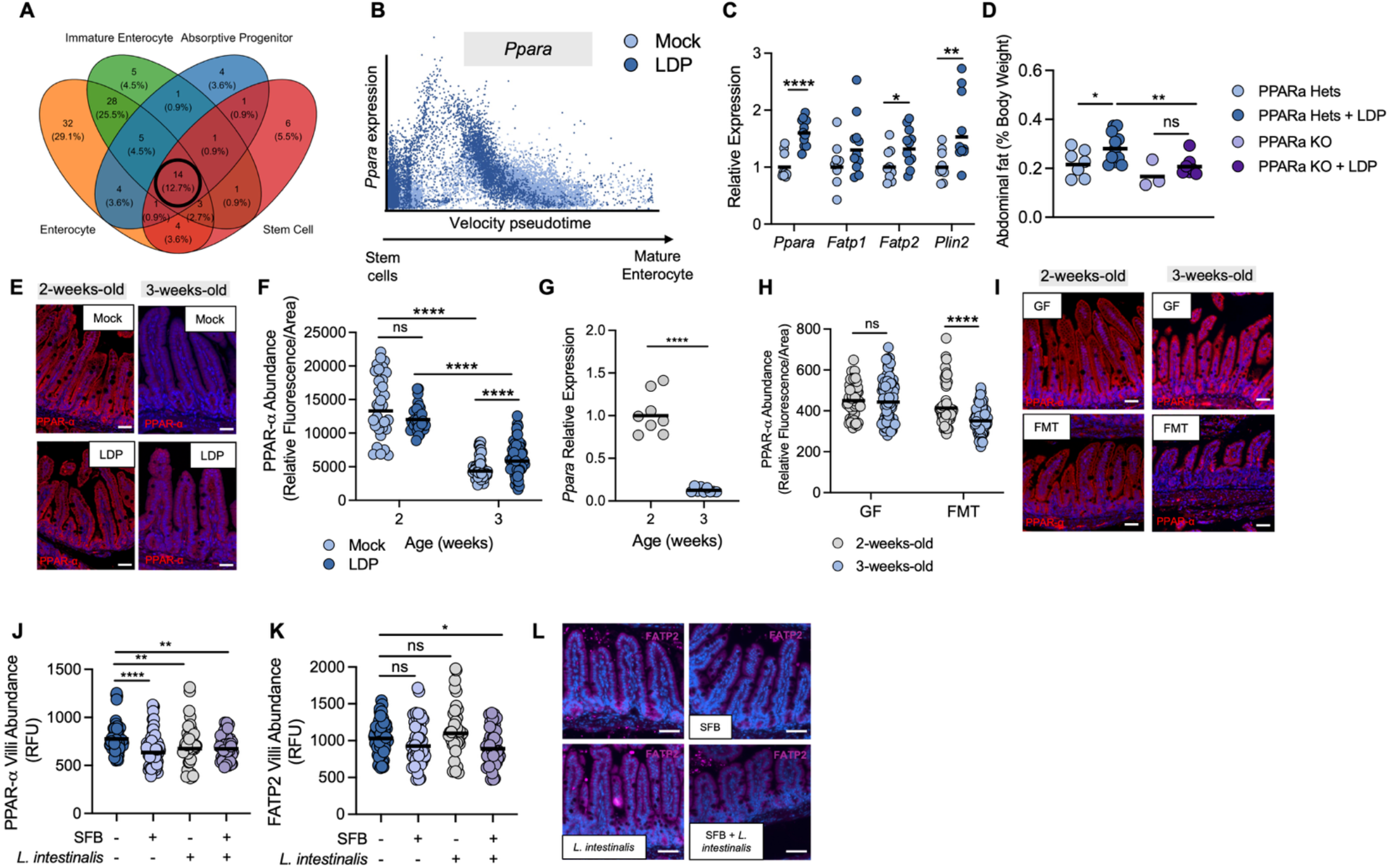
Early-life microbiota suppresses intestinal PPARα to reduce excess adiposity. (**A**) Venn diagram of genes upregulated in LDP absorptive epithelial cells across different cell types and supported by differential transcriptional kinetics inferred by RNA velocity analysis. Black circle indicates genes upregulated across all cell types with *Ppara* included in this group. (**B**) Latent time plot of showing *Ppara* expression along the differentiation trajectory of absorptive cells based upon scRNA-seq. (**C**) Expression of *Ppara* and its’ target genes in bulk epithelial cells isolated from the ileum epithelium of 3-week-old Mock and LDP-treated mice measured by qPCR. (**D**) Abdominal fat (% total body weight) of littermate PPARα knockout (KO) and heterozygotes exposed to LDP for the first three weeks of life or left untreated. (**E**) Representative images from immunofluorescent staining of PPAR-α in the ileum of indicated groups. (**F**) Quantification of PPARα abundance in the villi of 2 and 3-week-old Mock and LDP-treated mice. (**G**) Expression of *Ppara* in bulk epithelial cells isolated from the ileum epithelium of 2-week-old and 3-week-old mock treated mice measured by qPCR. (**H**) Quantification of PPAR-α abundance (by fluorescent intensity) in the villi of germ-free (GF) or conventionalized 2 and 3-week-old mice. **(I**) Representative images from immunofluorescent staining of PPARα in the ileum of indicated groups in **H. (J and K**) Quantification of (**J**) PPARα and (**K**) FATP2 abundance (by fluorescent intensity) in the villi of 3-week-old LDP-treated mice given SFB, *L. intestinalis*, or both. (**L**) Representative images from immunofluorescent staining of FATP2 in the ileum of the indicated groups in **J**. (C, D, G) Each dot represents one mouse. (F, H, J, and K) Each dot represents one villi. Data based on images from n = 5 – 7 mice. (C and G) *, p < 0.05; **, p < 0.01; ****, p < 0.0001 using an unpaired Student’s t test. (F and H) ****, p < 0.0001 using a two-way Anova with Šídák’s multiple comparisons test. (J and K) *, p < 0.05; **, p < 0.01; ****, p < 0.0001 using a one-way Anova with Dunnett’s multiple comparisons test. (D) *, p < 0.05; **, p < 0.01 using an unpaired Welch’s t test.

We determined that the SI microbiota expands during 2 to 3 weeks of life and that this development protects mice against antibiotic-induced fat accumulation (**Fig. 1A** and **Fig. 3, I and J**). As we hypothesized that SI microbiota development regulates intestinal *Ppara* and that disruption of the SI microbiota by early-life antibiotics drives PPARα-dependent increases in adiposity, we next asked how PPARα abundance in the SI changed between 2 and 3 weeks of life. In 2-week-old mice, PPARα was highly abundant in the SI, and no difference between mock and LDP-treated mice was observed (**Fig. 4, E and F**). By 3 weeks of age, both PPARα abundance and *Ppara* expression had significantly decreased in distal SI epithelial cells but remained significantly higher in LDP-treated compared to mock-treated mice (**Fig. 4, C, E–G**). Importantly, this reduction in PPARα abundance was microbiota-dependent, as intestinal PPARα levels remained equally elevated in GF 2- and 3-week-old mice (**Fig. 4, H and I**). As SFB and *L. intestinalis* were the two microbes significantly increased in the SI microbiota of 3-week-old mice (**Fig. 1, E and F**), we next examined intestinal PPARα abundance in antibiotic-exposed mice following administration of SFB, *L. intestinalis*, or both microbes (**Fig. 3K** and **Fig. 4J**). SFB and *L. intestinalis* were both able to reduce PPARα abundance in LDP-treated mice (**Fig. 4J**). However, only the combination of microbes reduced the abundance of the PPARα target gene FATP2 (**Fig. 4, K and L**). We show that intestinal PPARα is suppressed by expansion of the early-life microbiota, and that antibiotic treatment results in elevated intestinal PPARα and PPARα-dependent increases in adiposity during infancy.

### Development of the early-life small intestine microbiota induces short-term production of IL-22, reducing PPARα, and protecting against excess adiposity

Having determined that intestinal PPARα is downregulated by the SI microbiota during early life, we next investigated how the SI microbiota communicates with IECs to decrease PPARα. Previous work has shown that microbiota metabolites can regulate intestinal PPARs (*41-43*). We therefore asked whether a metabolite in the SI content of mock 3-week-old mice could suppress PPARα in enteroids from LDP-treated mice. However, supernatants from both the SI content of mock and LDP-exposed mice increased *Ppara* expression in enteroids (**Fig. 5A**), suggesting an alternative mechanism by which the microbiota suppresses PPARα. The activity and abundance of PPARα are inhibited by pro-inflammatory cytokines (*44, 45*), which raised the possibility that expansion of the SI microbiota during early life prompted an immune reaction that downregulated PPARα. We performed multiplex cytokine profiling of ileum tissue homogenate from 3-week-old mock and LDP-treated animals. Antibiotic treatment significantly reduced multiple cytokines, including IL-17A and TNFα (known regulators of intestinal lipid metabolism), in the ileum (**Fig. S5A**) (*46, 47*). We also measured levels of IL-22, another cytokine known to modulate intestinal lipid uptake (*48, 49*), in the ileum tissue homogenate and observed a strong reduction in LDP-treated mice (**Fig. 5B**). As various cytokines were decreased due to early-life antibiotics (**Fig. 5B** and **Fig. S5A**), we performed an initial screen in a fetal intestinal epithelial cell line (HIEC-6) to investigate which cytokine impacted PPARα activity in IECs (**Fig. 5C** and **Fig. S5B**). IL-22 was the only cytokine that reduced the expression of *FATP1*, a target gene of PPARα, *in vitro* (**Fig. 5C**). In subsequent experiments in enteroids from 3-week-old mock-treated mice, IL-22 significantly decreased *Ppara* expression (**Fig. 5D**). Furthermore, IL-22 reduced lipid uptake in HIEC-6 cells treated with the PPARα agonist WY14643 (**Fig. 5, E and F**), consistent with downregulation of PPARα-specific lipid uptake (*40*).

**Figure 5.**
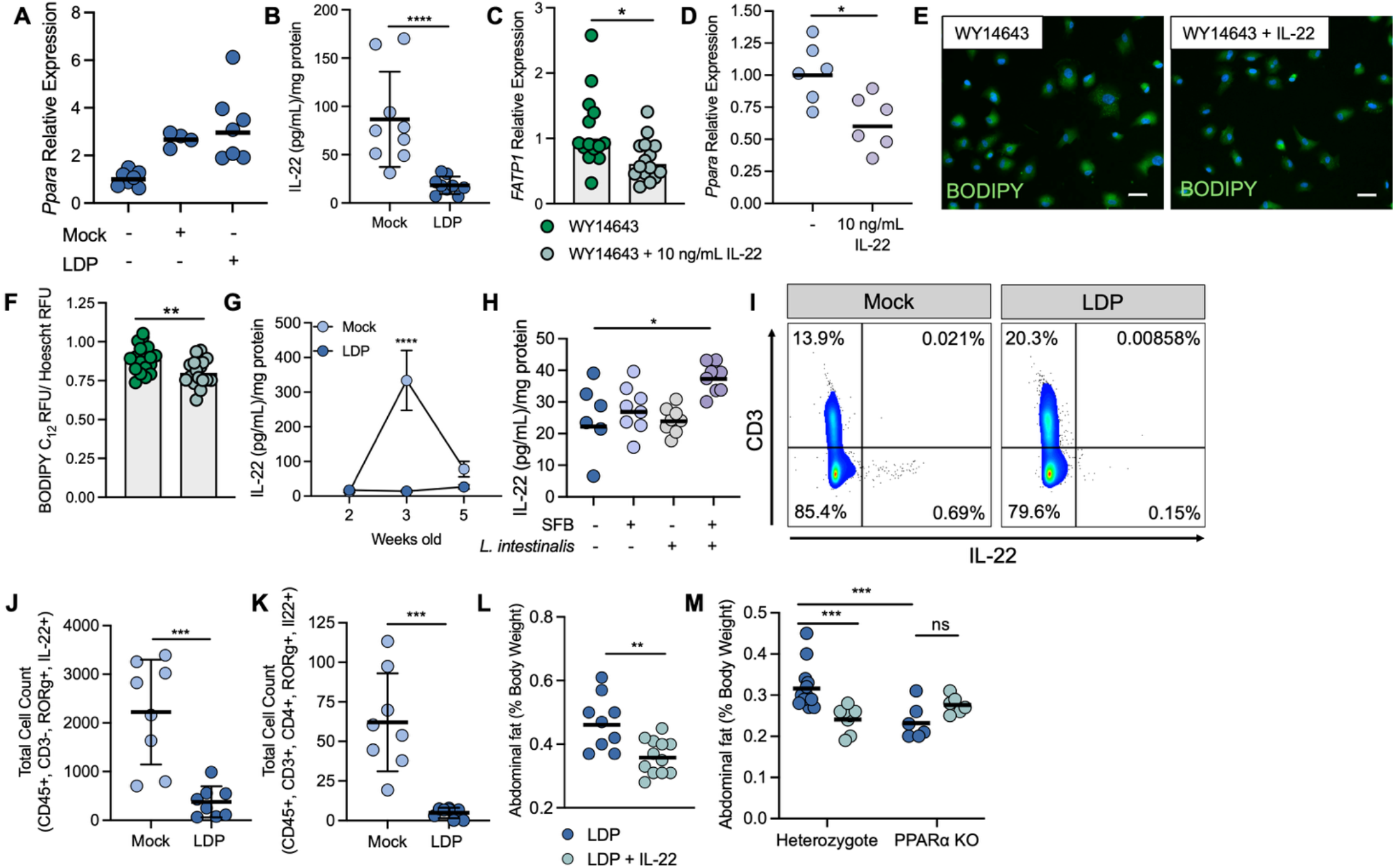
Early-life microbiota induces IL-22 production to suppress PPARα signaling and protect against adiposity. (**A**) Relative expression of *Ppara* in enteroids from 3-week-old LDP-treated mice cultured with supernatants from Mock or LDP small intestinal content. (**B**) Concentration of IL-22 in ileum tissue homogenate from 3-week-old mock and antibiotic-treated mice as determined by ELISA. (**C, E and F**) Human fetal epithelial cells (HIEC-6) were pre-treated with a PPAR-α agonist (WY14643) before incubation with IL-22 for 6 hours. Then, (**C**) RNA was extracted and expression of *FATP1* was measured by qPCR. (**D**) Expression of *Ppara* in enteroids isolated from 3-week-old mock mice. Enteroids were either mock treated or given 10 ng/mL IL-22 for 48 hours. **(E and F**) After treated as described above, HIEC-6 cells were then given 2.5 µM BODIPY-C12 for 1.5 hours before fixation. (**E**) Representative immunofluorescence images of BODIPY-C12 uptake. (**F**) BODIPY-C12 fluorescent intensity was then measured and normalized to Hescht nuclear fluorescence intensity. (**G**) Concentration of IL-22 in the ileum tissue homogenate of 2, 3, and 5-week-old Mock and LDP-treated mice as determined by ELISA. (**H**) Concentration of IL-22 in the ileum tissue homogenate in 3-week-old antibiotic treated mice that were orally gavage when they were 16, 18, and 20 days with either PBS, SFB, *L. intestinalis*, and both SFB and *L. intestinalis*. (**I - K**) Lamina propria was isolated from the ileum of mock and antibiotic-treated mice and enriched for CD45+ cells. Cells were then stimulated for 4 hours and stained. (**I**) Representative flow cytometry staining of the lamina propria from 3-week-old mock and antibiotic treated mice. (**J**) IL-22 positive cells in CD45+, CD3-, RORg+ population. (**K**) IL-22 positive cells in CD45+, CD3+, CD4+, and RORg+ population. (**L**) Abdominal fat (% total body weight) of LDP-treated mice treated with 1 µg of IL-22 (or mock treated) when mice were 15, 18, and 21 days old. (**M**) Abdominal fat (% total body weight) of littermate PPARα knockout (KO) and heterozygote mice exposed to LDP and treated with 1 µg of IL-22 (or mock treated) when mice were 15, 18, and 21 days old. (A and D) Each dot represents a technical replicate. Experiment was performed two independent times (n = 3). (B, H, J – M) Each dot represents one mouse. (C and D) Each dot represents a technical replicate. Experiments were performed three independent times. (F) Each dot represents the mean ± standard error of the mean (n = 6 – 8 mouse/group). (B – D, F, J – L) *, p < 0.05; **, p < 0.01; ***, p < 0.001; ****, p < 0.0001 using an unpaired two-tailed Student’s t test. (F) ****, p < 0.0001 (comparing Mock and LDP IL-22 at 3-weeks-old, 2-week-old and 3-week-old Mock IL-22, and 3-week-old and 5-week-old Mock IL-22) using a two-way Anova with Šídák’s multiple comparisons test. (H) *, p < 0.05 using a one-way Anova with Dunnett’s multiple comparisons test. (M) **\*\*\***, p < 0.001 using a two-way Anova with Šídák’s multiple comparisons test.

To understand if the early-life SI microbiota regulates IL-22 production, we measured IL-22 in the ileum of 2-, 3-, and 5-week-old mock and LDP-treated mice (antibiotic treatment was stopped at 3 weeks for 5-week-old LDP-exposed mice) (**Fig. 5G**). Intestinal IL-22 levels peaked at 3 weeks of age in mock-treated mice, and IL-22 abundance was significantly higher in the SI of 3-week-old mice than in 2- and 5-week-old mice (**Fig. 5G**). This induction of IL-22 was not observed in the ileum of antibiotic-treated or young GF mice, nor was there a difference in IL-22 in the colon of mock and antibiotic-exposed young mice (**Fig. 5G and Fig. S6, A and B**). The dynamics and localization of IL-22 production are consistent with previous reports that the SI microbiota elicit a weaning response specifically in the ileum (*33*). Consistently, we did not observe a significant difference in IL-22 levels in adult mice treated with LDP for the same period as young mice (**Fig. 3G** and **Fig. S6C**), indicating that antibiotics’ ability to prevent IL-22 induction is unique to early life.

Our results demonstrated that IL-22 is induced by the expansion of the SI microbiota that occurs between 2 and 3 weeks of life. Therefore, we asked whether reintroducing SFB, *L. intestinalis*, or both microbes to the SI during early life was sufficient to increase IL-22 levels in antibiotic-treated mice (**Fig. 5H**). LDP-treated mice that received both SFB and *L. intestinalis* had a significant increase in IL-22 compared to antibiotic-exposed mice that were mock-treated (**Fig. 5H**). Importantly, gavage with SFB or *L. intestinalis* alone was insufficient to significantly increase SI IL-22 concentrations (**Fig. 5H**).

As IL-22 is produced by both Th17 and group 3 innate lymphoid cells (ILC3s) (*50, 51*), we next investigated which cell types were responsible for the increase in SI IL-22 levels observed in 3-week-old mock-treated mice. We isolated the SI lamina propria from 3-week-old mock and antibiotic-exposed mice and enriched for CD45+ cells. Then, cells were stimulated and stained for IL-22 and markers of various immune cell types.

The majority of IL-22-positive cells were CD3^-^ (**Fig. 5, I–K**), but significant reductions in IL-22 in LDP-treated mice were observed in both CD3-negative and CD3-positive populations (**Fig. 5, J and K**). To investigate whether CD3^+^ cells were responsible for IL-22 production in our model, we treated control mice with an anti-CD3 antibody during 2 to 3 weeks of life. Treatment with anti-CD3 did not decrease IL-22 levels in 3-week-old mice, suggesting that CD3^+^ cells are not the source of IL-22 during weaning (**Fig. S7D**). Instead, we observed a significant increase in IL-22 levels in 3-week-old mice given anti-CD3 (**Fig. S7D**). This finding is consistent with previous work that revealed T regulatory (Tregs) cells are induced at the end of the weaning reaction, leading to decreased cytokine production (*33*). Therefore, the depletion of CD3^+^ cells in mock mice may be reducing Tregs, resulting in the observed elevation in IL-22 levels (**Fig. S7D**). Nonetheless, the increased IL-22 levels in mice treated with an anti-CD3 antibody suggest that ILC3s are the main producers of IL-22 in our early-life model, a finding supported by previous examination of ILC3 activation during weaning (*52*).

As the increased adiposity in antibiotic-treated mice was dependent on PPARα (**Fig. 4D**), we next asked if IL-22-dependent downregulation of PPARα protected mice exposed to LDP from excess adiposity. To restore IL-22 levels in antibiotic-exposed young mice, we treated LDP-exposed wild-type (WT) mice (**Fig. 5L**) and, in a separate experiment, littermate PPARα heterozygous and PPARα KO mice (**Fig. 5M**) with IL-22 during the 2-to-3-week window when IL-22 peaks in mock-treated mice (**Fig. 5G**). We then measured abdominal fat at 3 weeks of age. IL-22 significantly reduced adiposity in male, LDP-treated WT mice (**Fig. 5L**). Significantly, IL-22 failed to reduce adiposity in LDP-treated PPARα KO male mice but did decrease fat accumulation in their heterozygous littermates (**Fig. 5M**). Together, we have identified a previously unexplored mechanism by which the early-life SI microbiota induces IL-22 to regulate lipid uptake in IECs by suppressing intestinal PPARα, thereby decreasing adiposity and explaining the correlations observed between infant antibiotic treatment and increased risk of childhood obesity.

## DISCUSSION

Multiple studies have shown correlations between antibiotic use during the first two years of life and an increased risk for childhood obesity (*7, 10, 36*). Disruption to the infant gut microbiota is proposed as a mechanism by which early-life antibiotics promote metabolic dysfunction (*17, 53*). However, it remains unknown what antibiotic-induced changes to the infant gut microbiota perturb host metabolism, leading to an increased risk for obesity. Due to the relative ease in sample collection, investigations into the early-life gut microbiota have largely characterized the fecal microbiota (*12, 13*). As the fecal microbiota composition is distinct from the microbiota in other regions of the gastrointestinal tract (*19*), the missing link between early-life antibiotics and increased risk for childhood obesity may be found in upper regions of the gastrointestinal tract, such as the small intestine. We investigated the development of the early-life SI microbiota and how it interacts with the host to regulate metabolism. Expansion of the SI microbiota, specifically SFB and *L. intestinalis*, triggers production of IL-22. IL-22, in turn, suppresses PPARα in the SI epithelium and protects against excess fat accumulation. We identified that perturbation to the development of the early-life SI microbiota disrupts epithelial-immune crosstalk that occurs during weaning, resulting in long-lasting increases in adiposity.

Here, we examined the development of the distal SI microbiota, noting significant differences between the early-life SI and the colonic microbiota (**Fig. 1, A and B**, and **Fig. S1D**). We determined that the bacterial abundance in the distal SI bloomed during 2 to 3 weeks of age in mice. In contrast, the bacterial abundance in the colon was similar during this period (**Fig. 1A**), revealing that findings on the development of the infant gut microbiota based on fecal samples are not necessarily conserved across regions of the gastrointestinal tract (*54, 55*). Due to difficulties in collecting samples from the small intestine, our understanding of the members of the early-life SI microbiota is based upon data from malnourished or sick infants and children (*56-58*). Therefore, mouse models allow us the opportunity to investigate the dynamics of SI microbiota development and identify potentially important members. This study focused on two early-life SI microbiota members, SFB and *L. intestinalis*, that significantly expanded in the distal SI from 2 to 3 weeks of life. SFB has largely been hypothesized to be a mouse-specific microbe, with few reports identifying this microbe in humans (*59, 60*). However, a recent publication identified a human SFB strain that possessed a similar filamentous pattern and nucleotide similarity to SFB that is seen in mice (*61*). In humans, SFB colonization peaks in early life and is in low abundance in the microbiota of adults, which supports our investigation into SFB’s role in host metabolism during early life (*61*). Additionally, *Lactobacillus* is also more abundant in the early-life human gut microbiota and the SI, with *L. intestinalis* specifically having been previously shown to have an important role in modulating intestinal epithelial function (*54, 62, 63*). Therefore, our model allowed us to investigate the function of two SI microbiota members that are relevant to early life in humans.

In this work, we observed that both SFB and *L. intestinalis*, either separately or together, could suppress intestinal PPARα and reduce antibiotic-induced adiposity (**Fig. 4J**). However, we determined that both microbes were needed to decrease the abundance of FATP2, a fatty acid transporter shown to be regulated by PPARα (*40*) (**Fig. 4, K and L**). Consistently, treatment with both SFB and *L. intestinalis* was necessary to increase IL-22 in antibiotic-treated mice (**Fig. 5H**). Thus, IL-22 may be needed to reduce PPARα transcriptional activity and protect against adiposity, but SFB and *L. intestinalis* may have additional, distinct PPARα independent effects on the host that protect against adiposity (**Fig. 3, K – M**). The mechanism by which both SFB and *L. intestinalis* are needed to increase IL-22 remains unknown, and future work will examine whether these microbes impact each other’s colonization, or whether SFB and *L. intestinalis* may act as a first and second signal necessary for IL-22 production (*64, 65*).

At weaning, the ileum microbiota increases in abundance, triggering a “weaning reaction” that results in a substantial immune response. Previous work carefully dissected this phenomenon and determined that this weaning reaction must occur at a specific time point (i.e., weaning) to properly train the immune system (*33, 66-68*). Inhibition of the weaning reaction via antibiotics or depletion of immune cells led to increased susceptibility to colitis and intestinal inflammation (*33, 66*). In addition to immune system education, the weaning reaction also promotes microbiota-dependent changes in intestinal stem cells, resulting in long-lasting changes to epithelial cell activity, which alter the risk of inflammatory diseases (*66*). Here, we propose that another function of the weaning reaction is to signal to the host to downregulate lipid uptake as mice transition from a milk-based diet to a chow diet. Past investigations into the weaning reaction focused on microbiota-mediated production of TNFα and IFNγ (*33*), but in this paper, we show that IL-22 is induced at weaning in 3-week-old mice and then decreases in the ileum as mice age. In contrast to previous work, we investigated how this peak of IL-22 regulates lipid uptake by interacting with PPARα and thereby protects against excess fat accumulation. Similar to the long-lasting impact of the weaning reaction observed in previously published work (*33, 66*), we show that mice treated with antibiotics continued to accumulate more fat than untreated mice, despite no longer receiving antibiotics, but the mechanism behind this sustained effect remains unknown. Importantly, adult mice treated with antibiotics do not gain more fat than untreated mice, consistent with the theory of the weaning reaction in that phenotypes are observed only when mice are perturbed during this developmental window (*33*). In this work, we revealed another facet of the weaning reaction and showed that, in addition to protection against intestinal inflammation, the weaning reaction is also critical in establishing metabolic health.

While intestinal PPARα has been shown to potentiate metabolic dysfunction by increasing lipid absorption and storage during consumption of an HF diet in adult mice (*40*), the activity of intestinal PPARα during early life is understudied. We determined that intestinal PPARα decreases in abundance between 2 and 3 weeks of life and that this downregulation is dependent on the SI microbiota, as no difference in intestinal PPARα is observed in 2- and 3-week-old GF mice (**Fig. 4, E – G**). Recent publications identified that microbiota metabolites can regulate intestinal PPARα (*41-43*), but here we show metabolite-independent regulation of PPARα by the early-life microbiota. We identified that IL-22 decreases intestinal PPARα activity *in vitro*, which we hypothesize decreases lipid uptake and thus protects against additional fat accumulation (**Fig. 5, C – F, L and M)**. Past work in adult mice revealed an interaction between IL-22 and *Ppara* and showed that IL-22 treatment decreased intestinal *Ppara* expression during consumption of a HF diet (*48*). We have expanded upon this work and revealed that specific members of the early-life SI microbiota induce IL-22 production during weaning, leading to decreased intestinal PPARα and a reduction in adiposity. In this work, we have identified a novel mechanism by which the development of the SI microbiota promotes metabolic health by regulating epithelial-immune crosstalk, providing new insight into the correlations between early-life antibiotic use and an increased risk for obesity.

## ACKNOWLEDGMENTS

We acknowledge the Translational Pathology Shared Resource supported by NCI/NIH Cancer Center Support Grant P30CA068485 and the Shared Instrumentation Grant S10 OD023475-01A1. The indirect calorimetry study was performed by the Vanderbilt Mouse Metabolic Phenotyping Center (DK135073, DK020593, and 1S10RR028101-01). Flow Cytometry experiments were performed in the VUMC Flow Cytometry Shared Resource. The VUMC Flow Cytometry Shared Resource is supported by the Vanderbilt Ingram Cancer Center (P30 CA68485) and the Vanderbilt Digestive Disease Research Center (DK058404). Immunofluorescent images were collected through the use of the Vanderbilt Cell Imaging Shared Resource (supported by NIH grants CA68485, DK20593, DK58404, DK59637, and EY08126) using a Zeiss LSM880 Airyscan Confocal Microscope (supported by NIH grant 1 S10 OD021630 1). C.D.S. was supported by the NIH (T32DK007673), Howard Hughes Medical Institute (HHMI), a 2025 Early Career Research Grant from The Obesity Society, and the Vanderbilt Institute for Clinical and Translational Research (VICTR VR73501). J.O. was supported by the NIH (T32AI095202-15). J.L. was supported by the NIH (T32AI112541). D.A. was supported by the NIH (T32DK007673). M.A.R.V.’s lab was supported by São Paulo Research Foundation (FAPESP; 2023/14946-5), National Council for Scientific and Technological Development (CNPq) and Coordenação de Aperfeiçoamento de Pessoal de Nível Superior - Brasil (CAPES) - Finance Code 001. V.D.N. (2021/00393-9) received fellowship from FAPESP (2021/00393-9). M.X.B. is a HHMI Freeman Hrabowski Scholar. Work in M.X.B.’s lab was funded by the NIH (R01DK131104-01) and The Pew Charitable Trusts (2022-A-19568). Graphic abstract generated with Biorender: de Brito, C. (2026) https://BioRender.com/4406kxf

## AUTHOR CONTRIBUTIONS

M.X.B. and C.D.S. designed and conceived the study. C.D.S., J.L., M.W.R., and M.N.J. performed all experiments. C.B.B., J.O., D.T.C., D.A., and M.N.J assisted with experimental design and data collection. V.D.N. and M.A.R.V performed scRNA sequencing analysis. N.K. analyzed metagenomic sequencing samples. M.N.J. and L.L analyzed food consumption, body composition, and indirect calorimetry results. All authors contributed to the data analysis and preparation of the manuscript. C.D.S. and M.X.B. wrote the manuscript, and all authors reviewed it.

## MATERIAL & METHODS

### Lead contact

Further information and requests for resources and reagents should be directed to and will be fulfilled by the lead contact, Mariana Byndloss.

### Materials availability

All unique/stable reagents generated in this study are available from the lead contact without restriction.

### Data availability

Metagenomic sequencing data has been deposited in the Sequence Read Archive (SRA) database under the accession number: PRJNA1464670. ScRNA-sequencing files have been uploaded to Gene Expression Omnibus under the accession number: GSE333176. This paper does not report original code. Any additional information required to reanalyze the data reported in this paper is available from the lead contact upon request.

### Experimental model details Animal Experiments

All experiments in this study were approved by the Institutional Animal Care and Use Committee (IACUC) at Vanderbilt University Medical Center. At the end of the experiment, mice were humanely euthanized by CO_2_ administration. Afterwards, mice were weighed and ileum (distal small intestine) content, ileum, colon, liver, or adipose tissue (for experiments described below), and samples for histopathology and immunofluorescence were collected. Epididymal (abdominal) fat from mice was removed and weighed.

#### Conventional mouse experimental details

Female and male C57BL/6N mice (#027), 8-week-old, were obtained from Charles River Laboratory. Male mice were single-housed for 1 week before breeding pairs were created. 18 days after breeder pairs were established, pregnant females were divided into two groups. One group was maintained on regular drinking water, while the second group was given low doses of penicillin (LDP) (Thermo Scientific #J62442.14, 6.67 mg/mL) in their drinking water. LDP was continually given in the drinking water until offspring were 3 weeks old. Litters that were smaller than 5 mice or larger than 10 mice were excluded from experiments. All breeders were given *ad libitum* access to chow. For some experiments, LDP was given in the drinking water only when offspring of the breeders were 2 weeks old until they were 3-weeks-old. At 3 weeks of age, offspring of breeders were either humanely euthanized or placed on a 60% fat diet (HF) (Research Diets Inc., #D12492) or a 10% fat control diet (LF) (Research Diets Inc., #D12450J) for 5 weeks. For long-term experiments, mice were weighed weekly.

To study the effect of antibiotics on adult mice, 8-week-old female and male C57BL/6N mice (#027) were obtained from Charles River Laboratory. The initial weight of the mice was collected, and then the mice were assigned to groups so that no significant differences in weight between groups occurred before beginning treatments. Groups of male and female mice were then given LDP in their drinking water or left untreated for 3 weeks. All mice were given *ad libitum* access to chow.

PPARα knockout (KO) mice (B6;129S4-*Ppara*^*tm1Gonz*^/J) (#008154) and wildtype (WT) C57BL/6J (#000664) (6 to 8 weeks old) were procured from The Jackson Laboratory. The microbiota of PPARα KO and WT mice was depleted using the following cocktail of antibiotics: Ampicillin sodium salt (Sigma-Aldrich #A9518), Metronidazole (Sigma-Aldrich #M3761), Neomycin sulfate hydrate (Thermo Scientific, #J61499.14), and Vancomycin HCl (Sigma-Aldrich #V2002). 2.5 milligrams of each antibiotic was given to each mouse every 24 hours for 5 days. After 5 days of antibiotic treatment, mice were inoculated by intragastric gavage with a fecal matter transplant (FMT) prepared from the distal small intestine, cecal, and colon content of 8-week-old female C57BL/6N mice (#027) from Charles River Laboratory. PPARα KO and WT mice were then bred together to generate PPARα heterozygote mice. The genotype of offspring was determined by PCR using protocol primers described by The Jackson Laboratory. Heterozygotes were then bred together in order to have WT, KO, and heterozygote offspring in each litter, or heterozygotes were bred to KO mice, with their offspring comprising of heterozygotes or KO mice. Pregnant mice were either left untreated or given LDP in the drinking water 18 days after breeder pairs were established. LDP was continuously given in the drinking water until the offspring of the breeder were 3 weeks old.

To modulate microbiota-immune interactions, mice were manipulated as follows between 2 and 3 weeks of age. LDP-exposed mice were inoculated by intragastric gavage with *Lactobacillus intestinalis* (1 x 10^9^ CFU in 0.1 mL MRS broth), segmented filamentous bacteria (SFB) FMT (0.1 mL, preparation detailed below), or both *L. intestinalis* and SFB (*L. intestinalis* pellet resuspended in SFB FMT such that each mouse received SFB and 1 x 10^9^ CFU in 0.1 mL) when they were 16, 18, and 20 days old. In a separate experiment, antibiotic-treated mice (from both Charles River and PPARα heterozygote breeders) were injected intraperitoneally with 1 μg of mouse recombinant IL-22 (BioLegend, #576204) when they were 15, 18, and 21 days old. To assess the role of CD3+ cells, 150 μg anti-CD3 (BioXCell, # BE0002) or an equal amount of the isotype control (BioXCell, #BE0090) was injected intraperitoneally into mock-treated mice when they were 15, 18, and 21 days old.

#### Gnotobiotic mouse experimental details

C57Bl/6N germ-free mice (Taconic, B6NTac – GF) (female and male, 8-week-old) were purchased from the National Gnotobiotic Rodent Resource center at the University of North Carolina at Chapel Hill and moved into a positive pressure cage system upon arrival. A subset of germ-free mice were colonized with an FMT prepared from Charles River mice (as described above). 18 days after breeder pairs were established, pregnant GF and FMT-colonized females were divided into two groups. One group was maintained on regular drinking water, while the second group was given low doses of penicillin (LDP) in their drinking water. LDP was continually given in the drinking water until the offspring were 3 weeks old.

### Bacterial culture conditions

*Lactobacillus intestinalis* was isolated from the ileum content of 3-week-old mock-treated C57BL/6N mice by plating the content on MRS agar plates (BD Difco # 288130). Colonies were then screened on MRS plates with 500 μg/mL vancomycin, with colonies susceptible to vancomycin being isolated for further characterization, as *L. intestinalis* was previously shown to be inhibited by vancomycin (*63*). DNA was extracted from isolated colonies using the PureLink Genomic DNA Mini Kit (Invitrogen #K1820-01) and species identified by 16s rRNA sequencing by Genewiz. *L. intestinalis* was routinely grown anaerobically at 37 °C in MRS broth (BD Difco # 288130) or on MRS agar plates. SFB was generously provided by Dr. Shipra Vaishnava and regularly cultivated by passaging fecal slurry into germ-free C57BL/6N mice (colony maintained by Byndloss Laboratory) via intragastric gavage.

### Culture of murine intestinal enteroids

3-week-old mice, either mock-treated or exposed to LDP, were humanely euthanized, and the ileum removed and opened. The ileum was placed into 5 mL ice-cold HBSS (Thermo Fisher #14175095). Ileums were cut into 2 mm pieces and added to fresh ice-cold HBSS. Samples were washed twice with ice-cold HBSS and then placed into 5 mL of chelation buffer (HBSS with 2 mM EDTA/EGTA). Samples were rocked in chelation buffer at 4 °C for 30 minutes. After incubation, tissue was allowed to settle, and the supernatant discarded. 10 mL HBSS was then added, and samples shaken horizontally for 1 minute (approximately 120 – 180 shakes). The resulting supernatant was poured through a 70 μm filter and stored on ice. This process was repeated 3 times. The flow-through was centrifuged at 200 x g for 5 minutes at 4 °C. Pellets were resuspended in 1 mL of HBSS and the number of crypts in each fraction was assessed. Fractions containing mostly crypts were combined and fractions containing single cells were discarded. Cell pellets were resuspended in Mouse Intesticult (Stemcell Technologies #06005 and Matrigel (Corning #356234) (final concentration of Matrigel was approximately 80%). 7 μL of the mixture was aliquoted into a 6-well non-tissue culture-treated plate (12-16 domes/ well) and incubated at 37°C until solidified. Media (Mouse Intesticult, 10 μM Y-27632 (bio-techne | TOCRIS #1254), and Normocin (InvivoGen #ant-nr-05)) was then added, and enteroids incubated at 37°C, 5% CO_2_. Clonogenicity was determined by measuring the ratio of crypts seeded to enteroids formed after 5 days of incubation. For experiments with IL-22 (BioLegend, #576204), crypts were cultured for 3 – 4 days (until enteroids formed) and then treated with 10 ng/mL IL-22 for 48 hours. Then, RNA was extracted and qPCR performed as described below.

### Cell lines

HIEC-6 cells (ATCC CRL-3266) were grown in HIEC-6 cell culture media (OptiMEM 1 Reduced Serum Medium (Gibco #31985070), 4% heat inactivated fetal bovine serum (FBS) (VWR #89510-188), 10 mM GlutaMax (Gibco #35050061), and 20 mM HEPES (Gibco #15630080)). Human EGF (Gibco #AF-100-15-500UG) was added directly to the tissue culture flasks with a final concentration of 10 ng/mL. Cells were passaged every 3-4 days in T75 flasks according to ATCC protocols.

### Method details

#### Indirect Calorimetry

A standard 12h light/dark cycle was maintained throughout the calorimetry studies. Mice were placed in metabolic cages located in the Mouse Metabolic Phenotyping Center at Vanderbilt University (RRID: SCIR_021939) in a temperature- and humidity-controlled dedicated housing room. Energy expenditure measures were obtained using a computer controlled indirect calorimetry system (Promethion, Sable Systems, Las Vegas, NV). The calorimetry system consists of 16 metabolic cages (identical to home cages with bedding) each equipped with water bottles and food hoppers connected to load cells for food and water intake monitoring, and all animals had ad libitum access to standard chow and water throughout the study. Due to the small size of the young mice, food intake was measured manually by placing a preweighed amount of food at the bottom of the cage and then measuring the amount eaten in a 24-hour period. The air within the cages is sampled through microperforated stainless steel sampling tubes located in the inner bottom rim of the cages to ensure that the cage air is sampled uniformly, and that the temperature inside the cages does not exceed the temperature within the room. Respiratory gases are measured with an integrated fuel cell oxygen analyzer, spectrophotometric CO2 analyzer, capacitive water vapor partial pressure analyzer and barometric pressure analyzer (GA3, Sable Systems, Las Vegas, NV). The system uses two GA-3 analyzers operating in parallel, devoted to eight cages apiece, to maximize throughput. Gas sensors are calibrated monthly with 100% N2 as zero reference and with a span gas containing known concentrations of CO2. The gain of the O2 channel is adjusted at each incurrent measurement so that incurrent O2, after correction for water vapor dilution and barometric pressure, yields a concentration of 20.94% STPD (standard temperature and pressure, dry). Promethion utilizes a pull-mode, negative pressure system. Two multi-channel mass flow generators measure and control air flows (FR8, Sable Systems, Las Vegas, NV). The excurrent flow rate is set at 2000 mL/min. Water vapor is continuously measured and its dilution effect on O2 and CO2 are mathematically compensated for in the analysis stream (*69*). Oxygen consumption and carbon dioxide (CO2) production are measured for each mouse at 5 min intervals for 30 seconds. Incurrent air reference values are determined after measuring every 4 cages. Respiratory quotient (RQ) is calculated as the ratio of CO2 production over O2 consumption. Energy expenditure is calculated using the Weir equation: Kcal/ hr=60*(0.003941*VO2+0.001106*VCO2) (*70*). Ambulatory activity is determined simultaneously every second with the collection of the calorimetry data. Ambulatory activity and position are detected with XYZ beam arrays (BXYZ-R, Sable Systems, Las Vegas, NV) with a beam spacing of 1.0 cm interpolated to a centroid resolution of 0.25 cm. Consecutive adjacent infrared beam breaks are counted and converted to distance, with a minimum movement threshold set at 1cm. Data acquisition and instrument control were coordinated by MetaScreen and the obtained raw data were processed using MacroInterpreter (Sable Systems, Las Vegas, NV) using an analysis script detailing all aspects of data transformation. The script is available on request from Sable Systems. Body composition was determined by NMR (Bruker Minispec).

#### Oral fat tolerance test

3-week-old mock and LDP-treated mice were weaned and then fasted for 4 hours. Then, mice were injected intraperitoneally with 0.5 g/kg tyloxapol (Sigma Aldrich #1704003; dissolved in sterile PBS) 30 minutes before they were given 100 μl of olive oil containing 20 μg of BODIPY 500/510 C_1_,C_12_ fatty acid (Invitrogen #D3823) via intragastric gavage. Mice were humanely euthanized 3 hours after gavage and blood was collected. Samples were then spun at 1000 x g for 20 minutes at 4 °C and the supernatant collected and stored at -80 °C. Serum samples collected from the oral fat tolerance test described above were diluted, and the amount of triglycerides was measured (Cell Biolabs STA-396). The distal small intestine was also harvested, the content removed, and the tissue swiss rolled. Swiss rolls were fixed in 4% paraformaldehyde (Electron Microscopy Sciences #15710-S) for 48 hours before being placed in 15% sucrose (Fisher Chemicals #S5-500) for 16 hours. Swiss rolls were then incubated in 30% sucrose for another 16 hours before being embedded in optimal cutting temperature (O.C.T.) compound (Fisher Scientific #23730571). 5 μm thick sections were cut and mounted to slides. Slides were stored at -80 °C until staining.

#### Quantification of Eubacteria abundance by qPCR

Content from the distal SI and colon was collected either 2 or 3-weeks after mice were born and frozen in liquid nitrogen. DNA was then extracted from the samples using the DNeasy PowerSoil Kit (Qiagen #47016) according to the manufacturer’s instructions. Analysis was performed as described previously (*71*). Bacterial DNA was normalized and used as a template for quantitative real-time PCR (qPCR) using iQ SYBR Green Supermix (Bio-Rad #1708882) or PowerUp SYBR Green Master Mix (Applied Biosystems #A25742) and primers listed in **Table S1**. Eubacteria abundance was determined by calculating the gene copy number for each sample using standard curves generated from the 16S rRNA gene of Eubacteria as previously described (*72*).

#### Metagenomic sequencing of SI and colon content

Content from the distal SI and colon was collected either 2 or 3-weeks after mice were born and frozen in liquid nitrogen. DNA was then extracted from the samples using the DNeasy PowerSoil Kit (Qiagen #47016) according to the manufacturer’s instructions. Sequencing libraries were prepared with Twist Library Preparation kits, and sequencing was performed on Illumina NovaSeq 6000, with 5 million reads obtained per sample according to the manufacturer’s instructions. FASTQ files were downloaded from VANTAGE’s online repository and quality checked with FastQC (v0.11.9) and MultiQC (v1.6). Quality trimming and adapter removal was done with BBDuk (v39.13) (Bushnell, B. BBMap – sourceforge.net/projects/bbmap/). Reads were then mapped to the mouse genome using bowtie2 (v2.5.4) and mapped reads were removed (*73*). The remaining SAM files were turned into BAM files, sorted, and converted into FASTQ format using SAMtools (v1.21) (*74*). Paired end reads were merged using FLASH (v1.2.11) (*75*). Taxonomic profiling was conducted with MetaPhlAn4, and functional profiling was performed with HUMAnN3 (*76*).

#### Histology

Tissue from the distal small intestine (either Swiss rolls or small sections) was fixed in 10% neutral buffered formalin, paraffin-embedded, and 5-μm-thick sections were cut and mounted on slides and stained with hematoxylin and eosin. Representative images were taken using a Leica DM750 microscope and a Leica ICC50W camera. Villi length and area were measured using ImageJ.

#### Immunofluorescent staining and analysis

For immunofluorescent staining, slides were deparaffinized in Histoclear (National Diagnostics #HS2021) or xylene (Fisher Chemicals #X54) and then rehydrated in descending ethanol concentrations (100%, 95%, 90%, 70%, 50%) for 5 minutes each. Slides were then washed in PBS three times before antigen retrieval. Antigen retrieval was performed by incubating slides in antigen retrieval buffer ((10 mM Tris; 0.5 mM EGTA; pH 9) for Alpi and UEA-1) or ((10 mM Sodium citrate, 0.05% Tween 20, pH 6.0) for PPARα and FATP2)) in a water bath (100 °C) for 15 minutes. Samples were then allowed to cool to room temperature on the bench for 30 minutes before being washed in PBS. For ALPI and UEA-1 staining, slides were blocked for one hour at room temperature in 10% donor rabbit serum. ALPI primary antibody (Santa Cruz Biotechnology # sc-271431) was added to slides (1:50 in 1% donor rabbit serum) and incubated overnight at 4 °C. Slides were then washed three times with PBS and then a secondary antibody was added (1: 1000 Goat anti-Mouse IgG, Alexa Fluor 546) (Thermo Fisher #A-11003) along with UEA-1 (1 μg/mL) (Millipore Sigma #L9006). For PPARα and FATP2 staining, slides were blocked for two hours in 3% bovine serum albumin, Fraction V (BSA) (Fisher Scientific #AAJ6578818). PPARα (Thermo Fisher #PA1822A) (1:200) and FATP2 (Proteintech #14048-1-AP) (1:500) primary antibodies were added to slides at the indicated dilutions and incubated overnight at 4 °C. Slides were then washed three times with PBS and then a secondary antibody was added (1:1000 Goat anti-Rabbit IgG (Invitrogen #A21245)). All slides were incubated with secondary antibodies for 1 hour at room temperature before being washed three times with PBS. Nuclei were then stained using Hoechst 33342 (0.1 μg/mL) (Thermo Fisher #62249) for ten minutes. Samples were then washed and coverslips mounted using ProLong Gold Antifade Mountant (Thermo Fisher #P36934).

For frozen tissue staining, slides were thawed at room temperature for 20 minutes and then rehydrated with PBS for 10 minutes (slides were prepared from mice given oral fat tolerance test as described above). Sections were permeabilized by incubating with 1% BSA + 0.05% saponin (Thermo Fisher #J63209.AK) for 10 minutes. Slides were then washed with PBS, and then the nuclei were then stained using Hoechst 33342 (0.1 μg/mL) (Thermo Fisher #62249) for ten minutes. Samples were then washed and coverslips mounted using ProLong Gold Antifade Mountant (Thermo Fisher #P36934).

For ALPI and UEA-1 staining, as well as PPARα staining in **Fig. 4, E and F**, imaging was performed on a Zeiss LSM880 Airyscan Confocal Microscope. Representative images shown in the paper were collected using a Nikon A1 Microscope equipped with 405, 488, 561, and 645 nm lasers and a Plan Fluor 40X/1.30 NA immersion objective. Images were contrast-enhanced and cropped using FIJI/ImageJ software (NIH). Abundance of ALPI was calculated using FIJI/ImageJ measurement tools. For the remaining PPARα, FATP2, and BODIPY 500/510 C_1_,C_12_ fatty acid slides, imaging was performed on a Nikon Eclipse Ti2 fluorescent microscope. Abundance of PPARα, FATP2, and BODIP 500/510 C_1_,C_12_ fatty acid was quantified using NIS-Elements. For imaging across all microscope modalities, acquisition parameters were matched between samples during image acquisition.

#### EdU Staining

5-ethynyl-2′-deoxyuridine (EdU) was prepared at 10 mM in PBS and 100 µL was injected i.p. per animal 90 min before euthanasia. Swiss rolls from the ileum of 3-week-old mock and LDP-treated mice were then fixed in 4% formalin followed by paraffin embedding. The slides were labeled with Click-It Plus EdU Kit (ThermoFisher Scientific #C10634) for incorporation of fluorescent azide (AlexaFluor 647), followed by counterstaining with Hoechst 33342 and then mounted with ProLong Gold Antifade Mountant (Thermo Fisher #P36934). Imaging was performed on a Zeiss LSM880 Airyscan Confocal Microscope.

#### Ki67 Staining

Slides were placed on either the Leica Bond Max or Bond-RX IHC stainer. All steps besides dehydration, clearing and coverslipping were performed on the Bond stainer. Slides were deparaffinized. Heat induced antigen retrieval was performed on the Bond Max using their Epitope Retrieval 2 solution for 20 minutes. Slides were placed in a Protein Block (Ref# x0909, DAKO, Carpinteria, CA) for 10 minutes. The sections were incubated with anti-Ki67 (Catalog #12202S, Cell Signaling Technology, Danvers, MA) diluted 1:1,000 for one hour. The Bond Refine Polymer detection system was used for visualization. Slides were then dehydrated, cleared and coverslipped. Representative images were taken using a Leica DM750 microscope and a Leica ICC50W camera.

#### Intestinal crypt isolation and preparation of single cell suspension

3-week-old mice, either mock-treated or exposed to LDP, were humanely euthanized and the ileum removed and opened. Two ileums were pooled together in 5 mL ice-cold HBSS (Thermo Fisher #14175095). Ileums were cut into 2 mm pieces and added to fresh ice-cold HBSS. Samples were washed twice with ice-cold HBSS and then placed into 4 mL of chelation buffer (HBSS with 2 mM EDTA/EGTA). Samples were rocked in chelation buffer at 4 °C for 30 minutes. After incubation, tissue was allowed to settle, and the supernatant discarded. 10 mL HBSS was then added, and samples shaken horizontally for 1 minute (approximately 120 – 180 shakes). The resulting supernatant was poured through a 70 μm filter and stored on ice. This process was repeated 3 times. The flow-through was centrifuged at 500 rpm for 5 minutes at 4 °C. Pellets were resuspended in 1 mL of HBSS and the number of crypts in each fraction was assessed. Fractions containing mostly crypts were combined and fractions containing single cells were discarded. Isolated crypts were washed in 10 mL HBSS + 0.4% BSA and resuspended in 50 μL HBSS. Crypts were then resuspended in 2 mL of digestion buffer (2.5 mg/mL DNAse (Thermo Fisher #18047019) and 5 mg/mL Protease (Millipore Sigma #P5380) in PBS) and incubated on ice until crypts dissociated into single cells (approximately 10 minutes) (*77*). Single cells were centrifuged for 5 minutes at 300 x g at 4 °C. Pellet was washed one time in HBSS before resuspending in 100 μL PBS + 2% FBS + 1 mM CaCl_2_ and then dead cells removed using the EasySep Dead Cell Removal (Annexin V) Kit (Stemcell Technologies #17899) according to the published protocol. The resulting cell population was counted and approximately 10,000 cells used for single cell RNA sequencing.

#### Single-cell RNA-sequencing, data processing, and analysis

Single cell RNA sequencing (scRNA-seq) was performed by VANTAGE, the genomic core laboratory at Vanderbilt University Medical Center. Sequencing was done using the 10X Genomics Chromium Controller. Libraries from single cell suspensions were prepared according to 10X Genomics protocol using Chromium Next GEM Single Cell 3’ GEM Library Kit. Library Quality Control analysis was performed using Qubit and BioAnalyzer to determine the concentration and size. Each library that passes the final QC undergoes qPCR using the KAPA library quantification kit and QuantStudio. Sequencing was then performed in paired-end mode using the Illumina NovaSeq 6000 and a read length of 150 base pairs.

Raw sequencing data were processed using Cell Ranger v6.0.1 (10x Genomics) with the 10x Genomics the *mm10* mouse reference to perfom alignment to transcriptome, cell barcodes assignment and Unique Molecular Identifiers (UMIs) counting. Gene expression count matrices were generated using the standard Cell Ranger pipeline with default parameters. Ambient RNA contamination was estimated and corrected using the SoupX package (version 1.6.2) by comparing raw and filtered expression matrices (*78*). Genes detected in fewer than three cells and cells expressing fewer than 200 genes were excluded from downstream analyses using Seurat v5.4.0 (*79*). Doublets were identified and removed using scDblFinder (version 1.16.0) (*80*), and cells exhibiting mitochondrial gene content greater than 25% were excluded (**Table S2**). Quality control was performed independently for each sample prior to dataset concatenation for integration.

Data were normalized using *LogNormalize* in Seurat prior to integration. To mitigate technical variability across samples, data integration was performed using the IntegrateData function in Seurat (*79*), incorporating sample identity as a covariate. Dimensionality reduction was performed by PCA, and the first 30 principal components were used for UMAP visualization and graph-based clustering. Cluster marker genes were identified using *FindAllMarkers* with the Wilcoxon rank-sum test and Bonferroni correction. Cell populations were manually annotated based on canonical marker genes for murine ileal epithelial populations and cluster-defining gene signatures (**Table S3**).

Differences in relative cell-type abundance between experimental groups were assessed using Speckle v0.0.3 (*81*). Differential gene expression analyses within specific cell types were performed in Seurat using FindMarkers, with significance defined as adjusted p-value < 0.05 and log2 fold-change > 0.25. Functional enrichment analyses were conducted using the clusterProfiler package (version 4.10.1) (*82*) to identify overrepresented Gene Ontology biological processes among significantly differentially expressed genes. Module scores were computed using the AddModuleScore function in Seurat (*83*). Absorptive function scores were calculated based on curated lists of nutrient transporter genes (*34*) **(Table S4)**. GSEA for PPARα related pathways were run using GSEA version 4.4 using custom gene sets **(Table S5)**.

##### RNA velocity and dynamic transcriptional modelling

For RNA velocity analyses, spliced and unspliced transcript counts were generated from the original FASTQ files using velocyto run10x (version 0.17) with alignment to the *mm10* reference genome (*84*). Loom files were generated independently for each sample. Only absorptive epithelial cell lineage were retained for velocity analyses. These subsets were extracted from the integrated Seurat object and converted to the AnnData format for downstream analyses.

RNA velocity analysis was performed using scVelo (version 0.2.5) (*85*). First- and second-order moments were calculated using scv.pp.moments. Gene-specific kinetic parameters were inferred using the dynamical model implemented in scv.tl.recover_dynamics, enabling estimation of kinetic parameters related to transcription, splicing, and degradation rates. RNA velocities were computed in dynamical mode using scv.tl.velocity. Reliably modeled genes were defined as velocity genes with a fit likelihood greater than 0.1. Cellular progression along transcriptional trajectories was quantified by computing latent time using scv.tl.latent_time. Genes with dynamic behaviour were ranked by fit likelihood, and the top 300 genes were selected for downstream analyses. Cluster-specific dynamical genes were identified using scv.tl.rank_dynamical_genes. Velocity confidence was assessed using scv.tl.velocity_confidence to evaluate the robustness of velocity estimates across cells. RNA velocity analyses were initially performed separately for each experimental condition, followed by dataset merging for comparative analyses. To identify condition-dependent transcriptional dynamics, velocity-associated genes were ranked using scv.tl.rank_velocity_genes, and a combined set of top-ranked genes from each condition was used as input for differential kinetic testing. Differential transcriptional kinetics between experimental conditions were assessed using scv.tl.differential_kinetic_test. To prioritize biologically relevant candidates, genes exhibiting differential transcriptional kinetics were intersected with genes identified as differentially expressed in absorptive epithelial cells. From this intersection, genes significantly upregulated in the antibiotic-treated condition relative to control were selected. This integrative approach yielded a final set of 14 genes, representing candidates supported by both altered transcriptional dynamics and increased steady-state expression.

#### Enterocyte isolation

3-week-old mice, either mock-treated or exposed to LDP, were humanely euthanized, and the ileum was removed and opened. The ileum was placed in 5 mL ice-cold HBSS (Thermo Fisher #14175095). Samples were then placed into 5 mL of chelation buffer (HBSS with 2 mM EDTA/EGTA). Samples were rocked in chelation buffer at 4 °C for 30 minutes. Then, samples were shaken vigorously for 1 minute, after which the tissue was removed and samples spun at 1000 x g for 5 minutes. The resulting pellet was snap frozen in liquid nitrogen and RNA was isolated as described below.

#### HIEC-6 experimental details

HIEC-6 cells were cultivated as described above. For gene expression analysis, HIEC-6 cells were seeded into 12-well plates at a concentration of 3 x 10^6^ cells/mL 48 hours before the experiment. Cells were pre-treated with 100 μm WY 14643 (Tocris #1312) for one hour and then media was removed, and cells were either treated with 100 μm WY 14643 or 100 μm WY 14643 and 10 ng/mL recombinant human IL-22 (R&D Systems #782-IL) for 6 hours. RNA was then isolated as described below. For measuring fatty acid uptake in HIEC-6 cells, cells were seeded into a black 96 well plate at a density of 0.5 x 10^5^ cells/mL and incubated overnight. Cells were then pre-treated with 100 μm WY 14643 (Tocris #1312) for one hour and then media was removed and cells were either treated with 100 μm WY 14643 or 100 μm WY 14643 and 10 ng/mL recombinant human IL-22 (R&D Systems #782-IL) for 6 hours. After 6 hours, media was removed and replaced with the same conditions plus 2.5 μm BODIPY 500/510 C_1_,C_12_ fatty acid (Invitrogen #D3823). Cells were incubated for 1 hour and 30 minutes before media containing BODIPY was removed. Cells were washed 2x with 0.1% BSA and then fixed with 2.5% paraformaldehyde. Cells were washed again and then the nuclei stained with 1 μg/mL Hoechst 33342. Fluorescence intensity of BODIPY and Hoechst 33342 was measured using the VANTAstar plate reader (BMG LABTECH). Representative images were taken with a Nikon Eclipse Ti2 fluorescent microscope.

#### RNA Isolation and quantitative real-time PCR

Enterocyte cell pellets, adipose tissue, and liver were homogenized using a FastPrep-24 and RNA isolated by the TRIzol method (Invitrogen #15596018) following the manufacturer’s protocol. RNA from HIEC-6 cells and enteroids was isolated using the RNeasy mini kit (Qiagen #74104) according to the kit’s instructions. RNA was reverse transcribed using iScript gDNA Clear cDNA Synthesis Kit (Bio-Rad #1725035). Quantitative real-time PCR was performed using iQ SYBR Green Supermix (Bio-Rad #1708882) or PowerUp SYBR Green Master Mix (Applied Biosystems #A25742) and the appropriate primer sets (**Table S1**). Relative gene expression was calculated using *Act2b* (mouse) and *18S* (human) as the housekeeping genes.

#### Cytokine measurements

At the indicated age, the distal small intestine or colon was harvested from each mouse and immediately snap frozen in liquid nitrogen. Tissue was then homogenized in TNE Buffer (Quality Biological #351302101) containing protease inhibitor (Sigma-Aldrich #P8340) (100 mg tissue/mL buffer) using an Omni TH Tissue Homogenizer. The tissue homogenate was then centrifuged at 1000 x g, and the soluble fraction was separated from the pelleted debris. The concentration of protein was then measured in the tissue homogenate using a Pierce BCA Protein Assay Kit (Thermo Scientific #A55864). Cytokines in Fig. S6 were measured by Eve Technologies using the Mouse High Sensitivity T-cell 18-Plex Discovery Assay Array (Eve Technologies #MDHSTC18) after all samples were adjusted to the same concentration (4.8 mg/mL). IL-22 was measured in the tissue homogenate using the Legend Max Mouse IL-22 ELISA Kit (BioLegend #436307) and normalized to the total protein analyzed.

#### Lamina propria isolation and flow cytometry

The distal small intestine of 3-week-old mock and LDP-treated mice was harvested, and Peyer’s patches were removed. Then, the tissue was opened and cut into small pieces. The tissue was incubated with HBSS and 10% FBS and 1 mM Dithiothreitol (DTT) (Thermo Scientific #R0861) and shaken at room temperature for 10 minutes. Samples were then shaken vigorously by hand for 30 seconds and then the tissue was removed. To remove epithelial cells, the tissue was placed into HBSS and 10% FBS and 5 mM EDTA and incubated at 37 °C for 10 minutes while shaking. The samples were then vigorously shaken by hand for 30 seconds and the tissue placed into a new tube of HBSS and 10% FBS and 5 mM EDTA. Samples were shaken at 37 °C for 10 minutes followed by 30 seconds of vigorous shaking by hand. Tissue was then incubated in digestion buffer (RPMI 1640 (Gibco #11875093) with 1 mg/mL collagenase D (Roche #11088858001), 100 μg/mL DNase I (Sigma-Aldrich #DN25), and 0.1 u/mL Dispase (Worthington Biochemical Corporation #NPRO) for 20 minutes at 37 °C and then homogenized using m_intestine_01 protocol with the gentleMACS Octo Dissociator. Homogenate was then passed through a 70 μm cell strainer. Cells were spun at 2000 rpm for 7 minutes, and the pellet resuspended in EasySep Buffer (Stemcell Technologies #20144). CD45-positive cells were then isolated using the EasySep Mouse CD45 Positive Selection Kit (Stemcell Technologies #18945) according to the manufacturer’s instructions. After enrichment for CD45+ cells, cells were stimulated with 1x cell activation cocktail (BioLegend #423301) and 1x BD GolgiPlug (Water Biosciences #555029) in T cell media (RPMI, 10%FBS, 1x Non-Essential Amino Acids (Gibco #11140050), 1 mM Sodium Pyruvate (Gibco #11360070), 24 mM HEPES (Gibco #11344041), 5.5 mM L-glutamine (Gibco #21051024), and 0.385 μM 2-Mercaptoethanol (Gibco #21985023)) for 4 hours. After activation, cells were stained with extracellular surface markers and Zombie Aqua (BioLegend #423101) (to stain dead cells) for 30 minutes at 4 °C. Cells were then fixed and permeabilized with BD Pharmingen Transcription Factor Buffer Set (Water Biosciences #562725) before being stained for cytokines and transcription factors of interest. Flow cytometric analysis was performed using a 5-Laser Fortessa (BD) with BD FASC Diva Software. Data analysis was done using FlowJo (BD Biosciences). Antibodies used for flow cytometry are listed in **Table S6**.

## QUANTIFICATION AND STATISTICAL ANALYSIS

Fold changes of ratios (mRNA relative expression) as well as percent abdominal fat, bacterial relative abundance, and relative fluorescence intensity were transformed logarithmically before statistical analysis. For quantification of ALPI intensity, ALPI intensity in the enterocytes lining each villus was measured by ImageJ and then background intensity was subtracted from each value. An unpaired Welch’s t-test was used on the transformed data to determine whether differences between groups were statistically significant (p < 0.05). Comparisons between more than 3 groups were determined using a one-way ANOVA followed by Tukey’s or Šídák’s multiple comparison test. When comparisons between multiple groups and containing two independent variables were assessed, statistically significant differences between groups were determined by two-way ANOVA followed by Tukey’s or Šídák’s multiple comparison test. Statistical analysis of scRNA-seq data is described in the methods section for scRNA-seq data analysis.

For statistical analysis of metagenomic data, Bray-Curtis Dissimilarities and PERMANOVA tests were conducted using the R package, vegan (v2.6-10). Differential abundance analysis was performed with Kruskal-Wallis tests from the coin package (v1.4-3), and p-values were adjusted using the Benjamini-Hochberg procedure in the stats package (v.4.4.1). PCoA plots and boxplots were generated with the ggplot2 package (v3.5.2).

