## Supplementary materials for "Small intestine microbiota development prevents early-life adiposity via IL-22-mediated intestinal PPARα suppression"

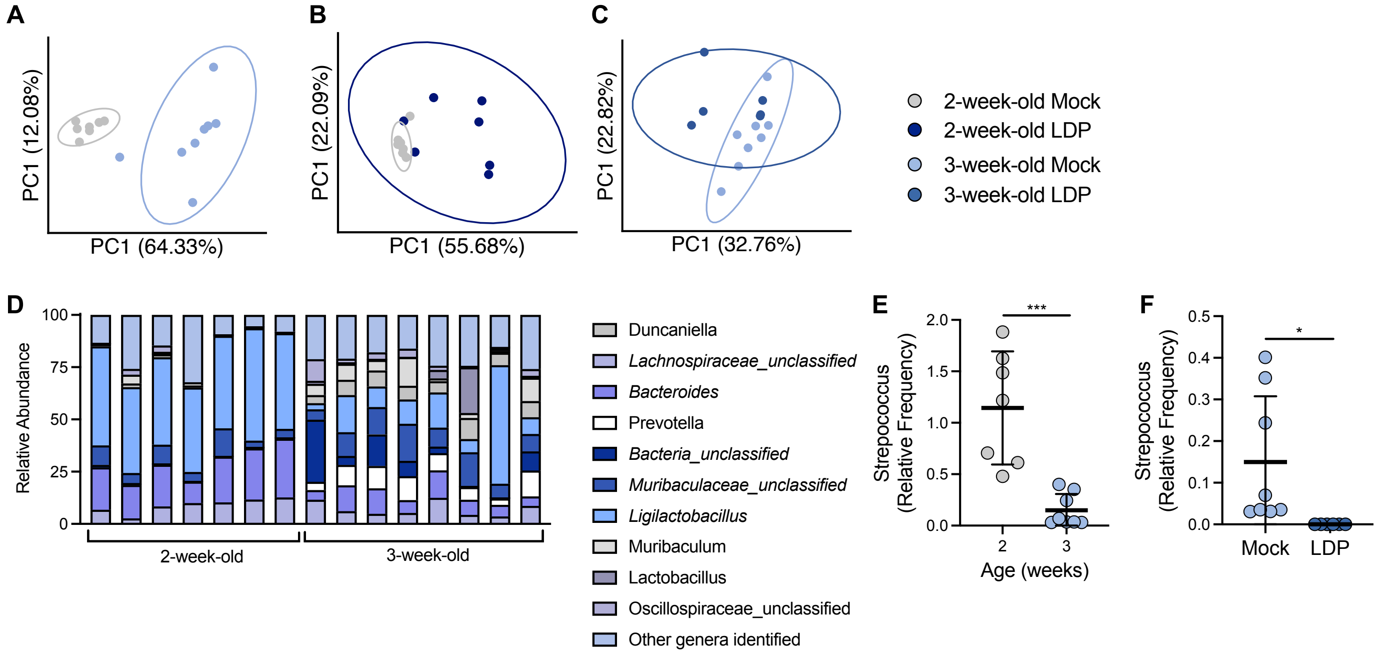


**Figure S1. Low-dose penicillin affects the 2-week-old colon microbiota. (A – C)** PCoA plot of genera in the colon content of (A) 2 and 3-week-old colon, (B) 2-week-old Mock and LDP, and (C) 3-week-old Mock and LDP based on Bray-curtis dissimilarity matrix. (D) Relative abundance of genera in the colon of mock mice at 2 and 3 weeks of age as determined by metagenomic sequencing. (E and F) Relative frequency of *Streptococcus* in (E) 2 and 3-week colon content and (F) 3-week-old Mock and LDP colon content. (E and F) *, $p < 0.05$; ***, $p < 0.001$ using an unpaired two-tailed Welch's t test.

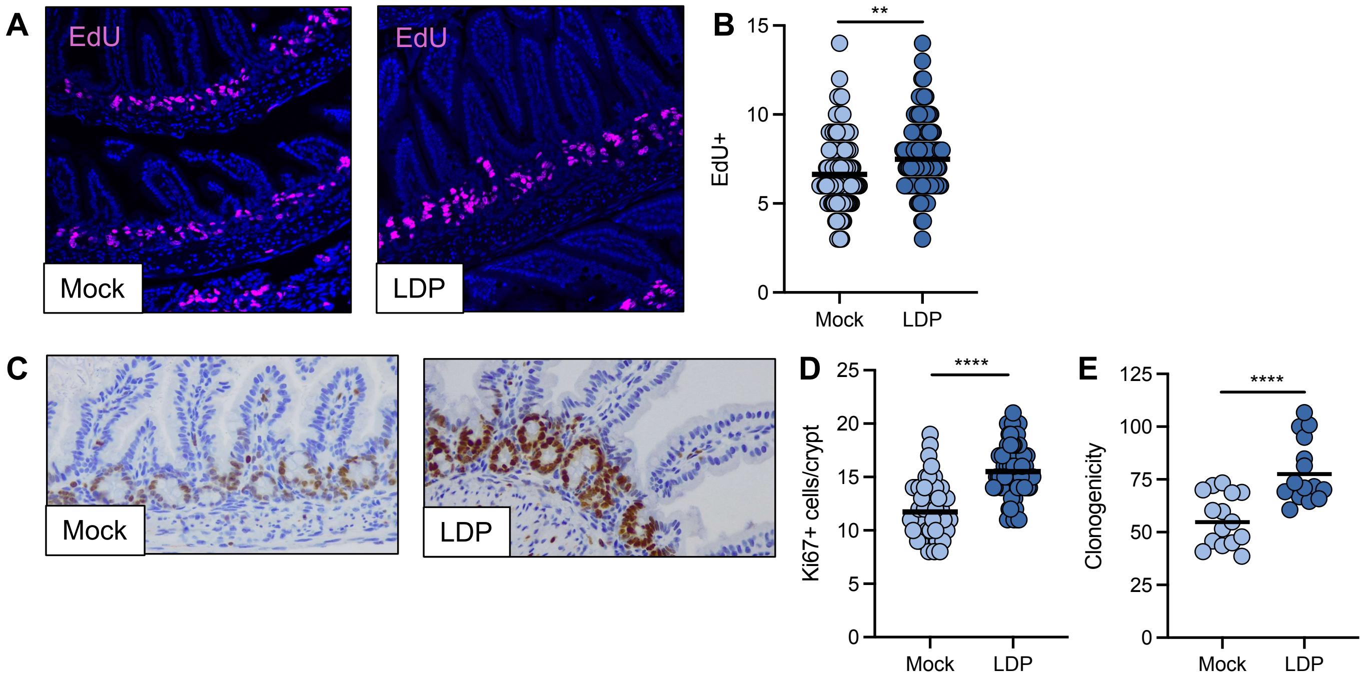


**Figure S2. Low dose penicillin (LDP) increases cell proliferation in the intestinal epithelium.** (A) Immunofluorescent imaging of EdU positive cells (magenta) in the ileum of 3-week-old Mock and LDP-exposed mice. (B) Quantification of EdU positive cells in the crypts of Mock and LDP-treated mice. (C) Representative images of Ki67 staining (brown) in 3-week-old ileum sections from each group. (D) Quantification of Ki67 positive cells in the crypts of Mock and antibiotic exposed mice. (E) Clonogenicity of crypts (number of enteroids formed/number of crypts seeded) isolated from the ileum of 3-week-old Mock and LDP treated mice. (B and D) Each dot represents one cell. Data representative of images from 6 – 7 mice. (E) Each dot represents one technical replicate (n = 3 technical replicates/mouse). Experiment performed 3 independent times. *, $p < 0.05$; **, $p < 0.01$; ****, $p < 0.0001$ using an unpaired two-tailed Student's t test.

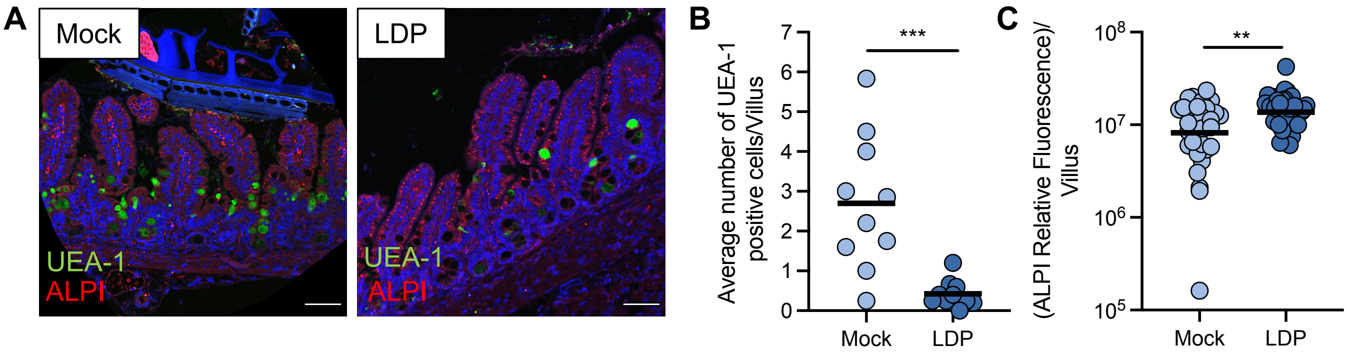


**Figure S3. Low dose penicillin (LDP) treatment disrupts differentiation of the intestinal epithelium.** (**A**) Representative immunofluorescent staining of Mock and LDP-exposed mice. Scale bar represents 100 μm (**B**) Quantification of UEA-1 positive cells in Mock and LDP-treated mice. Each dot represents the average UEA-1 positive cells per villus in one mouse. (**C**) Quantification of the relative fluorescent intensity of alkaline phosphatase (ALPI) in the villi of Mock and LDP mice. Each dot represents one villus. Data representative of images from six mice. (B and C) **, $p < 0.01$; ***, $p < 0.001$ using an unpaired Student's t test.

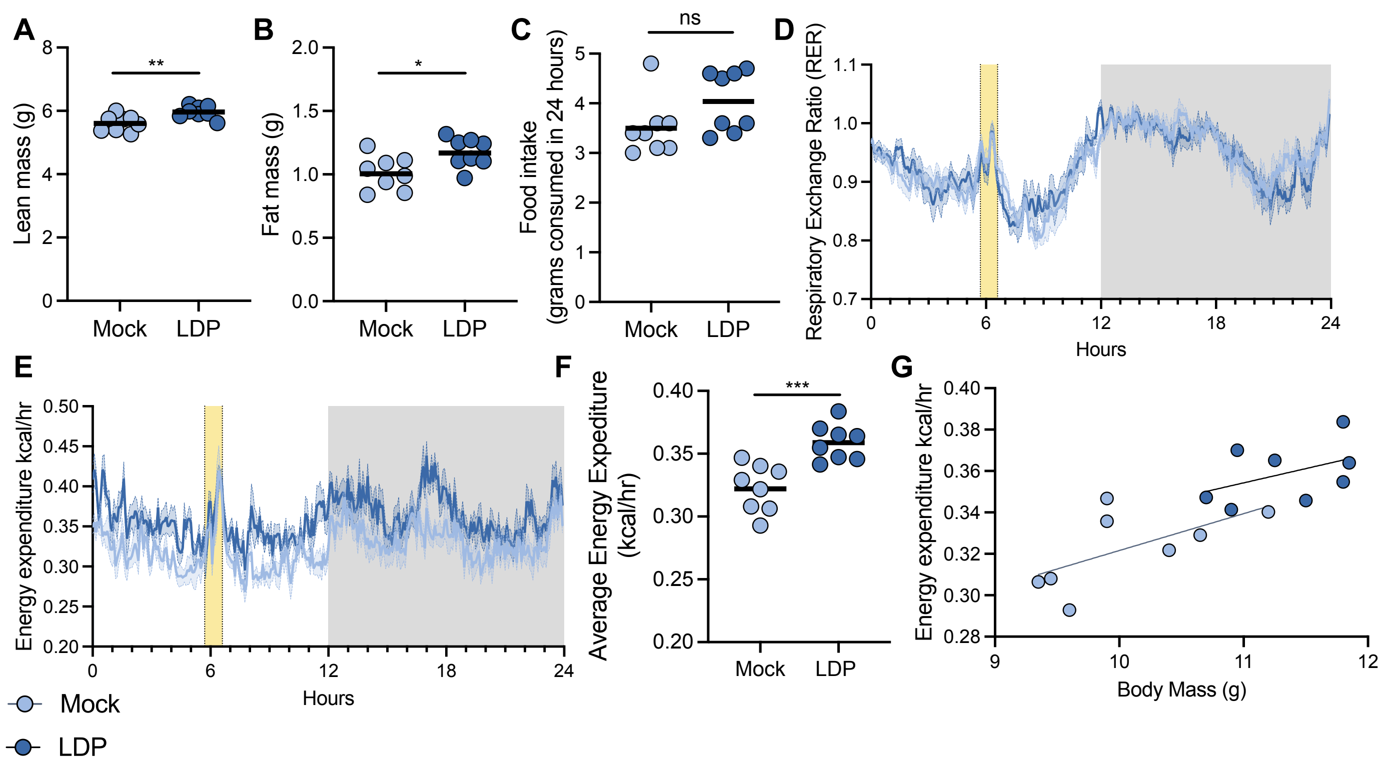


**Figure S4. Metabolic changes due to early-life antibiotic treatment.** (A and B) Body composition (lean mass (A) and fat mass (B) of 3-week-old Mock and LDP-treated mice was determined by NMR (Bruker Minispec). (C – G) 3-week-old Mock and LDP-treated mice were weaned, then placed in metabolic cages (1 mouse/cage) and allowed to acclimate for 3.5 days. Then, (C) food intake over 24 hours was measured manually. (D) Oxygen consumption and carbon dioxide production were used to calculate a Respiratory Exchange Ratio for each mouse over 24 hours. Grey rectangle indicates dark cycle. (E) Energy expenditure measures were obtained using indirect calorimetry over 24 hours (grey bar indicates dark cycle), and an average energy expenditure (F) was calculated for each mouse. (G) Average energy expenditure versus body mass. Each dot represents one mouse. (A – C, F) Line represents the geometric mean. *, $p < 0.05$; **, $p < 0.01$; ***, $p < 0.001$ using an unpaired two-tailed Student's t test. (D and E) Section in yellow: time where the cages were opened to weigh/dispense diet. This section of data was not included in the 12-hour averages of EE or RER, or in the ANCOVA stats. (G) ANCOVA statistics: 24-hour Energy Expenditure, total mass effect $p = 0.0308$, group effect $p = 0.1791$.

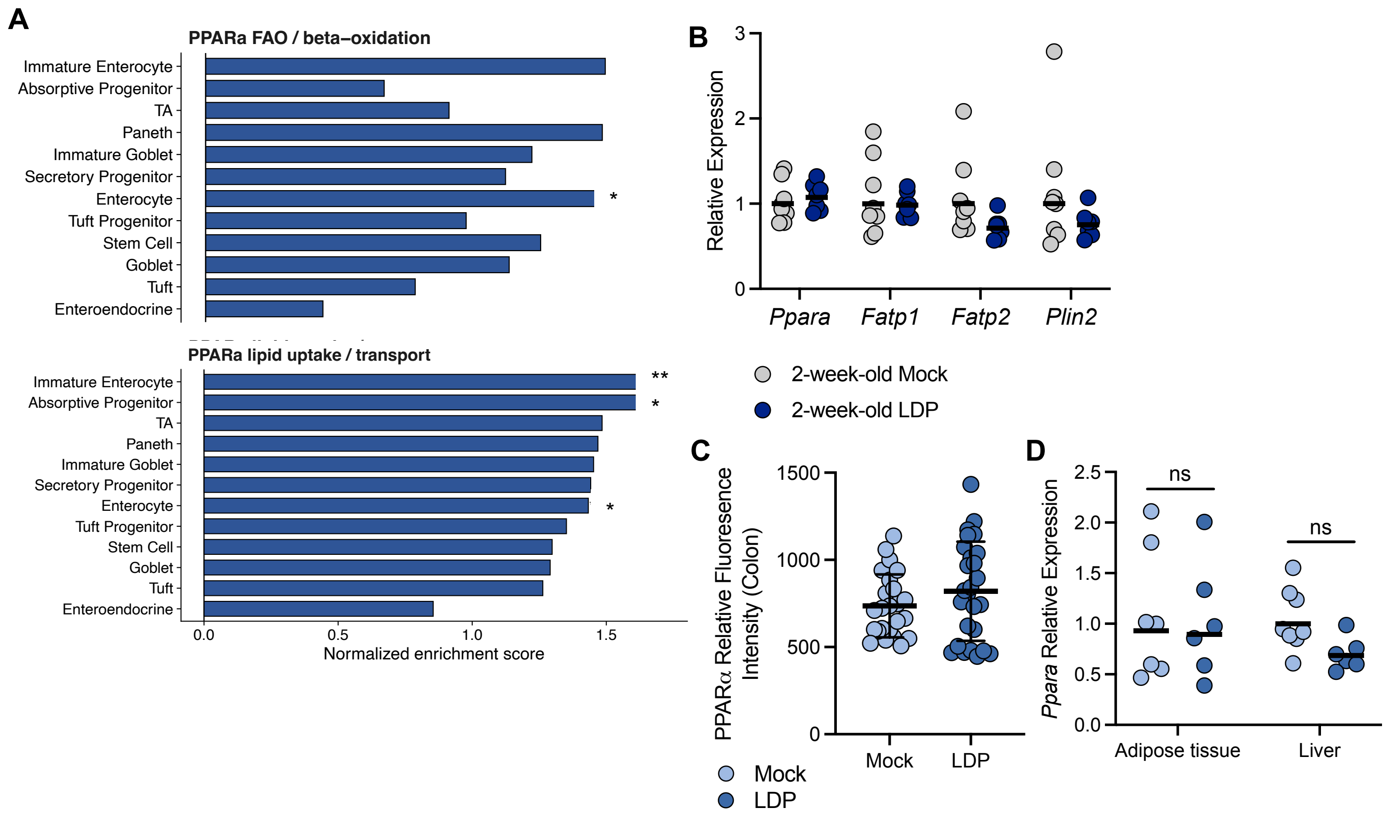


**Figure S5. Low dose penicillin (LDP) treatment specifically impacts *Ppara* expression in the 3-week-old ileum.** (A) Gene set enrichment analysis (GSEA) of indicated, upregulated PPARα related pathways in scRNAseq dataset from 3-week-old crypts from mock and antibiotic exposed mice. (B) Expression of Ppara and its' target genes in bulk epithelial cells isolated from the ileum epithelium of 2-week-old Mock and LDP-treated mice measured by qPCR. (C) Relative fluorescence intensity of PPARα in the colon of 3-week-old Mock and LDP-treated mice. (D) Relative expression of Ppara in the adipose tissue and liver of 3-week-old Mock and LDP-exposed mice.

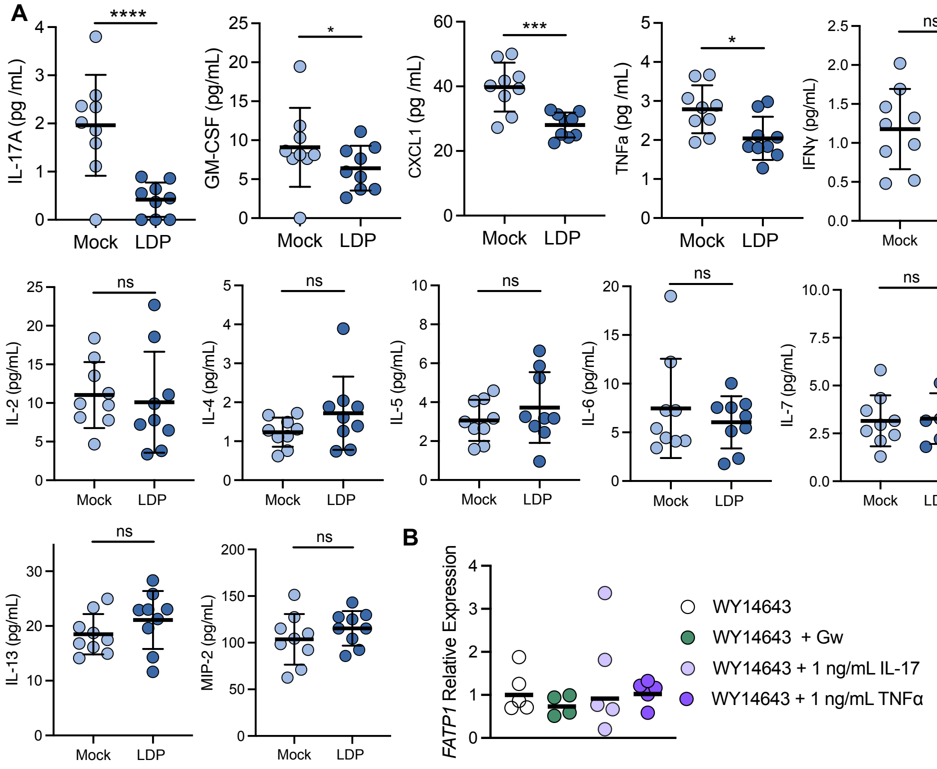


**Figure S6. Early-life antibiotics impacts intestinal immune profile, resulting in specific changes to PPARα activity. (A**) Level of indicated cytokines in tissue homogenate from the ileum of 3-week-old mock and antibiotic-treated mice as determined by Luminex multiplex assay. (**B**) Human fetal epithelial cells (HIEC-6) were pre-treated with a PPAR-α agonist (WY14 643) before incubation with IL-22 for 6 hours. Then, RNA was extracted and expression of *FATP1* was measured by qPCR. (A – D) Each dot represents one mouse. (E) Each dot represents one technical replicate. Experiment performed two independent times. (A – D) *, $p < 0.05$; ***, $p < 0.001$; ****, $p < 0.0001$ using an unpaired Student's t test.

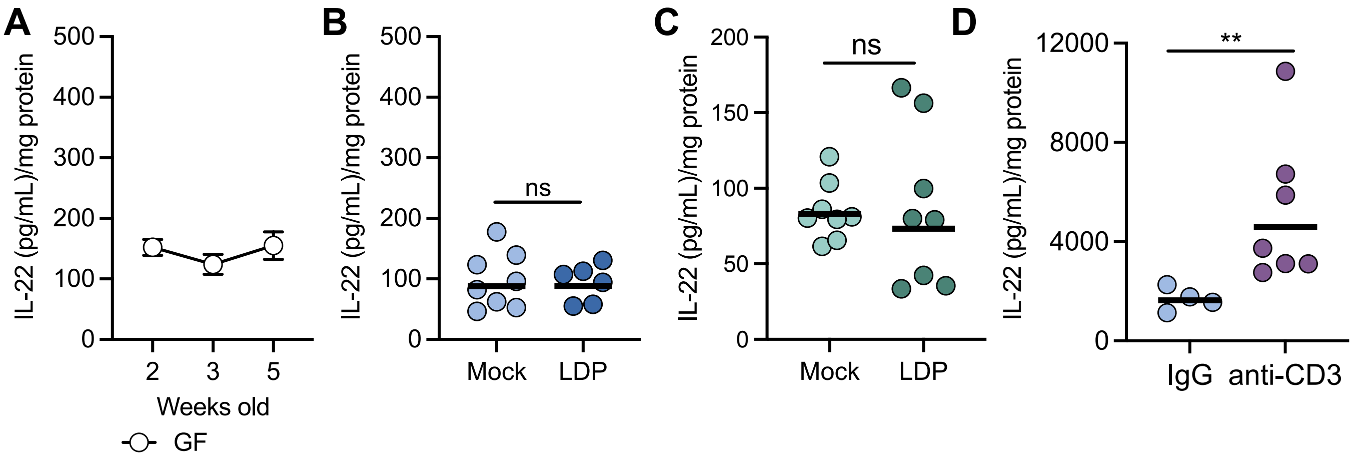


**Figure S7. Small intestine microbiota induces IL-22 production by CD3 negative cells. (A)** Concentration of IL-22 in the ileum tissue homogenate of 2, 3, and 5-week-old germ-free mice as determined by ELISA. (**B**) Concentration of IL-22 in colon tissue homogenate of 3-week-old Mock and LDP exposed mice. (**C**) Concentration of IL-22 in colon tissue homogenate of 11-week-old Mock and LDP exposed mice. Mice were mock treated or given antibiotics for 3 weeks, starting when they were 8-weeks-old. **(D**) Concentration of IL-22 in the ileum tissue homogenate of 3-week-old control and anti-CD3 treated mice. (A) Each dot represents the mean ± standard error of the mean (n = 5). (B – D) Each dot represents one mouse. (D) **, p < 0.01 using an unpaired Welch's t test.

**Table S1. Primers for RT-qPCR used in this study**

| RT-qPCR | | | |
|---|---|---|---|
| **Organism** | **Target gene** | **Forward primer (5' - 3')** | **Reverse primer (5' - 3')** |
| *Eubacteria* | *16S rRNA* | ACTCCTACGGGAGGCAGCAGT | ATTACCGCGGCTGCTGGC |
| *Mus musculus* | *Act2b* | GCTGAGAGGGAAATCGTGCGTG | CCAGGGAGGAAGAGGATGCGG |
| *Mus musculus* | *Ppara* | AGAGCCCCATCTGTCCTCTC | ACTGGTAGTCTGCAAAACCAAA |
| *Mus musculus* | *Fatp1* | CGCTTTCTGCGTATCGTCTG | GATGCACGGGATCGTGTCT |
| *Mus musculus* | *Fatp2* | GATGCCGTGTCCGTCTTTTAC | GACTTCAGACCTCCACGACTC |
| *Mus musculus* | *Plin2* | GACCTTGTGTCCTCCGCTTAT | CAACCGCAATTTGTGGCTC |
| *Homo sapiens* | *18S* | GATATGCTCATGTGGTGTTGA | ACGTTCCACCTCATCCTCA |
| *Homo sapiens* | *FATP1* | GGGGCAGTGTCTCATCTATGG | CCGATGTACTGAACCACCGT |

**Table S2. QC metrics for scRNAseq dataset**

| Sample | Condition | Estimated Number of Cells | Mean Reads per Cell | SoupX Score | scDblFinder Positive | Final Cell Number | Final Number of Genes |
|---|---|---|---|---|---|---|---|
| 11392-CS-0001 | *mock-treated* | *8,249* | *38,562* | *0.047* | *1687* | *6376* | *16618* |
| 11392-CS-0002 | *mock-treated* | *10,150* | *43,083* | *0.042* | *2160* | *7800* | *16974* |
| 11392-CS-0003 | *mock-treated* | *12,194* | *30,635* | *0.056* | *2411* | *9553* | *17231* |
| 11392-CS-0004 | *LDP-exposed* | *3,524* | *93,890* | *0.002* | *775* | *2735* | *15609* |
| 11392-CS-0005 | *LDP-exposed* | *14,026* | *26,768* | *0.034* | *2024* | *11732* | *17644* |
| 11392-CS-0006 | *LDP-exposed* | *7,053* | *61,113* | *0.037* | *1520* | *5404* | *16509* |

**Table S3. Marker genes used to identify cell type in scRNA-seq dataset**

| Cell Identity | Genes |
|---|---|
| Epithelial | Epcam |
| Epithelial | Krt8 |
| Epithelial | Krt18 |
| Endothelial | Pecam1 |
| Stromal | Col1a1 |
| Stromal | Col1a2 |
| Stromal | Vwf |
| Immune | Cd52 |
| Immune | Cd2 |
| Immune | Cd3d |
| Immune | Cd3g |
| Immune | Cd3e |
| Immune | Cd79a |
| Immune | Cd79b |
| Immune | Cd14 |
| Immune | Cd68 |
| Immune | Cd83 |
| Immune | Csf1r |
| Immune | Fcer1g |
| Immune | Ptprc |
| *Stem Cell* | *Lgr5* |
| *Stem Cell* | *Ascl2* |
| *Stem Cell* | *Smoc2* |
| *Stem Cell* | *Rgmb* |
| *Transit-Amplifying* | *Mki67* |
| *Transit-Amplifying* | *Top2a* |
| *Transit-Amplifying* | *Pcna* |
| *Paneth* | *Defa5* |
| *Paneth* | *Defa6* |
| *Paneth* | *Reg3a* |
| *Mature Goblet* | *Clca1* |
| *Goblet* | *Spdef* |
| *Goblet* | *Fcgbp* |
| *Goblet* | *Zg16* |
| *Goblet* | *Agr2* |
| *Goblet* | *Muc2* |
| *Enterocyte* | *Rbp2* |
| *Enterocyte* | *Anpep* |
| *Enterocyte* | *Fabp2* |
| *Enteroendocrine* | *Chga* |
| *Enteroendocrine* | *Chgb* |
| *Enteroendocrine* | *Neurod1* |
| *Tuft* | *Pou2f3* |

| | |
|---|---|
| *Tuft* | *Lrmp* |
| *Tuft* | *Trpm5* |
| *Tuft* | *Avil* |
| *M Cell* | *Spib* |
| *M Cell* | *Ccl20* |
| *M Cell* | *Gp2* |

**Table S4. Gene sets used to calculate nutrient absorption score**

| Nutrient | Gene |
|---|---|
| *LIPID* | *APOA1* |
| *LIPID* | *APOC3* |
| *LIPID* | *APOB* |
| *LIPID* | *FABP6* |
| *LIPID* | *FABP2* |
| *LIPID* | *PLIN2* |
| *LIPID* | *PLIN3* |
| *LIPID* | *SAR1B* |
| *LIPID* | *ABCG2* |
| *LIPID* | *SLC27A4* |
| *LIPID* | *APOM* |
| *LIPID* | *APOC2* |
| *LIPID* | *MGAT3* |
| *LIPID* | *LPL* |
| *LIPID* | *ACAT1* |
| *LIPID* | *ACSL3* |
| *LIPID* | *HMGCR* |
| *LIPID* | *ACAT2* |
| *LIPID* | *FFAR4* |
| *BILE SALT* | *SLC51B* |
| *BILE SALT* | *SLC51A* |
| *BILE SALT* | *SLC10A2* |
| *BILE SALT* | *SLC27A2* |
| *BILE SALT* | *SLC5A9* |
| *BILE SALT* | *ABCG5* |
| *VITAMIN* | *RBP2* |
| *VITAMIN* | *TCN2* |
| *VITAMIN* | *CYP4F2* |
| *VITAMIN* | *SLC23A1* |
| *VITAMIN* | *SLC52A1* |
| *VITAMIN* | *RDH5* |
| *VITAMIN* | *CYP4F3* |
| *VITAMIN* | *SLC23A3* |
| *VITAMIN* | *SLC22A4* |
| *VITAMIN* | *VNN1* |
| *VITAMIN* | *BTD* |
| *VITAMIN* | *BCO1* |
| *VITAMIN* | *BCO2* |
| *VITAMIN* | *CD320* |
| *VITAMIN* | *DRBP4* |
| *WATER* | *AQP1* |
| *WATER* | *AQP3* |
| *WATER* | *AQP7* |

| | |
|---|---|
| *WATER* | *AQP11* |
| *WATER* | *AQP8* |
| *AMINO ACID* | *SLC7A9* |
| *AMINO ACID* | *SLC6A19* |
| *AMINO ACID* | *SLC3A1* |
| *AMINO ACID* | *SLC7A7* |
| *AMINO ACID* | *SLC3A2* |
| *AMINO ACID* | *SLC1A1* |
| *AMINO ACID* | *SLC6A20* |
| *AMINO ACID* | *SLC6A6* |
| *AMINO ACID* | *SLC1A7* |
| *AMINO ACID* | *SLC25A39* |
| *AMINO ACID* | *SLC38A2* |
| *AMINO ACID* | *SLC38A1* |
| *AMINO ACID* | *SLC1A5* |
| *AMINO ACID* | *SLC25A13* |
| *AMINO ACID* | *SLC7A6* |
| *AMINO ACID* | *SLC38A5* |
| *AMINO ACID* | *SLC25A12* |
| *AMINO ACID* | *SLC1A4* |
| *AMINO ACID* | *SLC43A1* |
| *WATER* | *SLC5A1* |
| *WATER* | *SLC37A4* |
| *WATER* | *SLC2A5* |
| *WATER* | *SLC5A9* |
| *WATER* | *SLC2A2* |
| *WATER* | *SLC5A11* |
| *WATER* | *SLC50A1* |
| *WATER* | *SLC2A10* |
| *WATER* | *SLC2A10* |
| *INORGANIC SOLUTE* | *SLC25A3* |
| *INORGANIC SOLUTE* | *SSLC20A1* |
| *INORGANIC SOLUTE* | *SLC9A3R1* |
| *INORGANIC SOLUTE* | *SLC9A4A7* |
| *INORGANIC SOLUTE* | *SLC13A1* |
| *INORGANIC SOLUTE* | *SLC34A2* |
| *INORGANIC SOLUTE* | *SLC26A2* |
| *INORGANIC SOLUTE* | *SLC4A4* |
| *INORGANIC SOLUTE* | *CFTR* |
| *ORGANIC SOLUTE* | *SLC51B* |
| *ORGANIC SOLUTE* | *SLC10A2* |
| *ORGANIC SOLUTE* | *SLC13A2* |
| *ORGANIC SOLUTE* | *SLC51A* |
| *ORGANIC SOLUTE* | *SLC16A5* |
| *ORGANIC SOLUTE* | *SLC16A4* |

| | |
|---|---|
| ORGANIC SOLUTE | SLC44A1 |
| ORGANIC SOLUTE | SLC44A3 |
| ORGANIC SOLUTE | SLC16A1 |
| ORGANIC SOLUTE | SLC5A3 |
| ORGANIC SOLUTE | SCL2A13 |
| ORGANIC SOLUTE | SLC22A23 |
| ORGANIC SOLUTE | SLC16A3 |
| ORGANIC SOLUTE | SLC26A6 |
| ORGANIC SOLUTE | SLC16A9 |
| ORGANIC SOLUTE | SLCO3A1 |
| METAL ION | SLC39A4 |
| METAL ION | SLC25A37 |
| METAL ION | SLC39A5 |
| METAL ION | SLC31A1 |
| METAL ION | SLC31A2 |
| METAL ION | SLC30A4 |
| METAL ION | SLC39A7 |
| METAL ION | SLC39A8 |
| METAL ION | SLC30A9 |
| METAL ION | SLC39A3 |
| METAL ION | SLC39A9 |
| METAL ION | SLC30A7 |
| METAL ION | SLC9A2 |
| METAL ION | ATP2A3 |
| METAL ION | SCNN1A |
| METAL ION | SCNN1B |
| METAL ION | KCNS3 |
| METAL ION | SLC30A5 |
| METAL ION | SLC30A6 |
| METAL ION | SLC25A28 |
| METAL ION | KCTD14 |
| METAL ION | KCTD17 |
| NUCLEOTIDE | SLC28A1 |
| NUCLEOTIDE | SLC25A36 |
| NUCLEOTIDE | SLC35A3 |
| NUCLEOTIDE | NT5E |
| NUCLEOTIDE | SLC35A1 |
| NUCLEOTIDE | SLC35B3 |
| NUCLEOTIDE | SLC35B2 |
| NUCLEOTIDE | SLC25A33 |
| NUCLEOTIDE | SLC35D1 |
| NUCLEOTIDE | SLC17A5 |
| NUCLEOTIDE | SLC28A2 |
| NUCLEOTIDE | SLC29A1 |
| NUCLEOTIDE | SLC29A2 |

| | |
|---|---|
| *NUCLEOTIDE* | *SLC17A9* |

**Table S5. Genes used for PPARα pathway GSEA**

| Pathway | Gene |
|---|---|
| Mitochondrial fatty acid import and oxidation | *Cpt1a* |
| Mitochondrial fatty acid import and oxidation | *Cpt2* |
| Mitochondrial fatty acid import and oxidation | *Acadvl* |
| Mitochondrial fatty acid import and oxidation | *Acadl* |
| Mitochondrial fatty acid import and oxidation | *Acadm* |
| Mitochondrial fatty acid import and oxidation | *Acads* |
| Mitochondrial fatty acid import and oxidation | *Hadha* |
| Mitochondrial fatty acid import and oxidation | *Hadbh* |
| Mitochondrial fatty acid import and oxidation | *Hadha* |
| Mitochondrial fatty acid import and oxidation | *Decr1* |
| Peroxisomal fatty acid oxidation | Acox1 |
| Peroxisomal fatty acid oxidation | *Acox2* |
| Peroxisomal fatty acid oxidation | *Ehhadh* |
| Peroxisomal fatty acid oxidation | *Acaa1a* |
| Peroxisomal fatty acid oxidation | *Acaa1b* |
| Ketogenesis/ fasting-like lipid catabolism | *Hmgcs2* |
| Ketogenesis/ fasting-like lipid catabolism | *Bdh1* |
| PPARalpha-linked metabolic switch | *Pdk4* |
| Fatty acid uptake / membrane-associated transport | *Cd36* |
| Fatty acid uptake / membrane-associated transport | Slc27a1 |
| Fatty acid uptake / membrane-associated transport | *Slc27a2* |
| Fatty acid uptake / membrane-associated transport | *Slc27a4* |
| Intracellular fatty acid binding / intestinal lipid handling | *Fabp1* |
| Intracellular fatty acid binding / intestinal lipid handling | *Fabp2* |
| Intracellular fatty acid binding / intestinal lipid handling | *Fabp5* |
| Fatty acid activation | *Acsl1* |
| Fatty acid activation | *Acsl3* |
| Fatty acid activation | *Acsl4* |
| Fatty acid activation | *Acsl5* |
| Lipid droplet/lipid handling | *Plin2* |
| PPARalpha-responsive lipid handling | *Angptl4* |

**Table S6. Antibodies used for flow cytometry**

| Antibody | Source | Identifier |
|---|---|---|
| anti-CD45.2 | BioLegend | Cat #109821 |
| anti-CD4 | BioLegend | Cat #100565 |
| anti-CD3 | BD Pharmingen | Cat #561825 |
| anti-TCR beta | BioLegend | Cat #109219 |
| anti-TCR gamma/delta | BioLegend | Cat #118119 |
| anti-CD8 alpha | BioLegend | Cat #100743 |
| anti-IFN gamma | BD Horizon | Cat #612769 |
| anti-ROR gamma | Invitrogen | Cat #17-6981-82 |
| anti-IL-17 | Invitrogen | Cat #25717782 |
| anti-IL-22 | BioLegend | Cat #516404 |
| anti-Foxp3 | BD Pharmingen | Cat #567453 |
| anti-TNF alpha | BioLegend | Cat #506341 |
